# Assembly principles and stoichiometry of a complete human kinetochore module

**DOI:** 10.1101/2020.12.01.407130

**Authors:** Kai Walstein, Arsen Petrovic, Dongqing Pan, Birte Hagemeier, Dorothee Vogt, Ingrid Vetter, Andrea Musacchio

## Abstract

Centromeres are epigenetically determined chromosomal loci that seed kinetochore assembly to promote chromosome segregation during cell division. CENP-A, a centromere-specific histone H3 variant, establishes the foundations for centromere epigenetic memory and kinetochore assembly. It recruits the constitutive centromere-associated network (CCAN), which in turn assembles the microtubule-binding interface. How the specific organization of centromeric chromatin relates to kinetochore assembly and to centromere identity through cell division remains conjectural. Here, we break new ground by reconstituting a functional full-length version of CENP-C, the largest human CCAN subunit and a blueprint of kinetochore assembly. We show that full-length CENP-C, a dimer, binds stably to two nucleosomes, and permits further assembly of all other kinetochore subunits *in vitro* with relative ratios that closely match those of endogenous human kinetochores. Our results imply that human kinetochores emerge from clustering multiple copies of a fundamental module, and may have important implications for trans-generational inheritance of centromeric chromatin.

## Introduction

In addition to the hereditary information enshrined in the genetic code, chromosomes carry a wealth of additional epigenetic information that controls their structure and accessibility in time and space. Epigenetic information is usually implemented through covalent modifications of the DNA itself, or through modifications of the histones, proteins that pack DNA into nucleosomes, the fundamental organizational unit of eukaryotic genomes (Allis and Jenuwein, 2016). In addition to these well-studied mechanisms, a most striking and unconventional example of epigenetic determination subtends to the establishment and maintenance of centromere identity (Fukagawa and Earnshaw, 2014; McKinley and Cheeseman, 2016; Musacchio and Desai, 2017). Centromeres are unique chromatin loci of the eukaryotic chromosome. They seed the assembly of kinetochores, large protein complexes that establish a point of contact between chromosomes and spindle microtubules during mitosis and meiosis (Musacchio and Desai, 2017). The realization over 20 years ago that centromere identity, with few exceptions, is established in a DNA-sequence-independent manner, triggered a crucial conceptual leap in centromere studies and redirected the spotlight onto the molecular basis of centromere persistence through cell division (Karpen and Allshire, 1997).

A most characteristic and almost ubiquitous trait of centromeres is the enrichment of the histone H3-like centromeric protein A (CENP-A) (Earnshaw and Rothfield, 1985; Sullivan et al., 1994). Like H3, CENP-A forms a tight tetramer with H4, which additionally binds two H2A:H2B dimers to form an octameric nucleosome core particle (Black and Cleveland, 2011). The CENP-A nucleosome is surrounded by a group of 16 proteins collectively named the constitutive centromere associated network (CCAN) and organized in different stable sub-complexes (Foltz et al., 2006; Hinshaw and Harrison, 2019; Izuta et al., 2006; Obuse et al., 2004; Okada et al., 2006; Yan et al., 2019). Two CCAN subunits, CENP-N and CENP-C, interact directly with CENP-A (Carroll et al., 2010; Carroll et al., 2009; Kato et al., 2013). In addition, CENP-C, predicted to be predominantly intrinsically disordered, interacts with several other CCAN subunits to stabilize the overall globular assembly of CCAN (Hinshaw and Harrison, 2013; Hinshaw and Harrison, 2019; Klare et al., 2015; McKinley et al., 2015; Nagpal et al., 2015; Pentakota et al., 2017; Tanaka et al., 2009; Watanabe et al., 2019; Weir et al., 2016; Yan et al., 2019). CENP-C and another CCAN subunit, CENP-T, also provide a platform for the assembly of the kinetochore’s outer layer (Gascoigne et al., 2011; Przewloka et al., 2011; Screpanti et al., 2011). The latter comprises three further sub-complexes, known as the KNL1, MIS12, and NDC80 complexes, and collectively referred to as the KMN network (Cheeseman and Desai, 2008). Besides important regulatory functions, the KMN network, through its NDC80 complex (NDC80C), provides a site for binding spindle microtubules, an interaction that promotes the alignment and segregation of chromosomes to the daughter cells (Cheeseman et al., 2006; DeLuca et al., 2006).

Kinetochores across eukaryotes display significant differences in complexity. The *S. cerevisiae* kinetochore assembles around ~125 base pairs (bp) of a defined centromeric DNA sequence (and therefore not epigenetically) that wraps around a single CENP-A^Cse4^ nucleosome, and binds a single microtubule, for which reasons it is identified as a ‘point’ kinetochore. Conversely, kinetochores in the majority of eukaryotes are identified as ‘regional’ because they extend over much larger centromeric DNA segments (up to Mbp in humans), expose multiple CENP-A nucleosomes, and bind multiple microtubules (Musacchio and Desai, 2017). This extreme range of complexity raises the question whether regional kinetochores are built by patching together multiple copies of a “unit module” related to the point kinetochore of *S. cerevisiae*. This idea remains untested, but it is plausible and several observations support it. First, subunit composition and physical interactions appear to be largely conserved in point and regional kinetochores, as demonstrated by comprehensive biochemical reconstitutions and structural analyses (Hinshaw and Harrison, 2019; Klare et al., 2015; McKinley et al., 2015; Pekgoz Altunkaya et al., 2016; Pesenti et al., 2018; Weir et al., 2016; Yan et al., 2019). Second, the relative ratios of kinetochore subunits appear to be conserved in point and regional kinetochores, suggesting regional kinetochores emerge from the multiplication of a fundamental point module (Joglekar et al., 2009; Joglekar et al., 2008; Joglekar et al., 2006; Suzuki et al., 2015; Wan et al., 2009). Finally, the same relative subunit ratios observed at human kinetochores are also observed in partial reconstitutions of single, discrete particles of human kinetochores (Pesenti et al., 2018; Suzuki et al., 2015; Weir et al., 2016), once again suggesting that the larger assembly is created by convolution of individual modules.

From a reductionist perspective, this hypothesis raises the crucial question whether an individual module recapitulates some or even all fundamental properties of the regional structure, including promotion of bi-orientation and epigenetic specification and inheritance. Answering this question may have important technological implications, particularly towards the development of artificial chromosomes, but is technically and conceptually challenging. Despite very considerable progress in our understanding of the structural organization of the core protein complex surrounding the CENP-A nucleosome (Hinshaw and Harrison, 2019; Pesenti et al., 2018; Yan et al., 2019), there is persistent uncertainty on the organization of the continuous segment of centromeric chromatin underlying regional kinetochores. For example, even if CENP-A is greatly enriched within human centromeric chromatin in comparison to chromosome arms, it is surrounded and vastly outnumbered by histone H3 (Bodor et al., 2014). Various models for the organization of regional centromeric chromatin have been discussed (Fukagawa and Earnshaw, 2014), but conclusive evidence to support one or the other is missing. Furthermore, despite progress in the isolation of point kinetochores (Akiyoshi et al., 2010; Lang et al., 2018), there are no robust procedures for the isolation of individual modules from natural regional kinetochores. To address these limitations, we applied bottom-up approaches based on biochemical reconstitution of human kinetochore sub-structures of increasing complexity, followed by iterative assessments and validation of their functional and structural properties in comparison to those of real kinetochores, for instance with regard to the role of KMN multivalency in kinetochore-microtubule attachment (Huis In ’t Veld et al., 2016; Huis In ’t Veld et al., 2019; Volkov et al., 2018).

Here, we engineered for the first time a full-length, dimeric form of human CENP-C. This allowed us to dissect how oligomerization and multivalency of CENP-C reflects on kinetochore assembly and the underlying chromatin. We demonstrate that full-length CENP-C can bind concomitantly to two individual nucleosomes, or to two adjacent nucleosomes in a dinucleosome. We show that this organization supports the robust recruitment of all other kinetochore subunits with predicted stoichiometries, and we present a thorough dissection of the interactions that make this possible. Our studies represent an important step towards the reconstitution of a minimal regional kinetochore module.

## Results

### Reconstitution of full-length CENP-C (CENP-C^F^) by protein fusion

Human CENP-C is a 943-residue protein (Figure 1A). Its N-terminal region interacts with the MIS12 complex (MIS12C) (Gascoigne et al., 2011; Petrovic et al., 2016; Przewloka et al., 2011; Screpanti et al., 2011). The following PEST (Proline-Glutamic acid-Serine-Threonine)-rich region binds the CENP-HIKM and CENP-NL CCAN subcomplexes (Hinshaw and Harrison, 2013; Klare et al., 2015; McKinley et al., 2015; Nagpal et al., 2015; Pentakota et al., 2017). The central region encompasses a CENP-A-selective binding motif (indicated as ‘1’ in Figure 1A) that interacts with an exposed acidic patch on histones H2A and H2B as well as with the C-terminal tail of CENP-A (Carroll et al., 2010; Guse et al., 2011; Kato et al., 2013). It is required for robust kinetochore localization of CENP-C (Carroll et al., 2010; Guo et al., 2017; Heeger et al., 2005; Milks et al., 2009; Song et al., 2002; Tanaka et al., 2009). The constellation of N-terminal region, PEST, and central region spans the entire depth of the kinetochore from its outer domain until the centromere, and we have speculated that it provides spatial clues (a blueprint) for kinetochore assembly (Klare et al., 2015). Implicit in this analogy is that CENP-C promotes cooperative interactions of kinetochore subunits that enhance the overall stability of the inner kinetochore complex. Indeed, a fragment of CENP-C encompassing MIS12C binding site, PEST, and central region (CENP-C^1-544^) was sufficient to reconstitute a recombinant 26-subunit kinetochore particle with a CENP-A nucleosome core particle (CENP-A^NCP^) and the majority of CCAN and KMN subunits (Klare et al., 2015; Pesenti et al., 2018; Weir et al., 2016).

**Figure 1.**
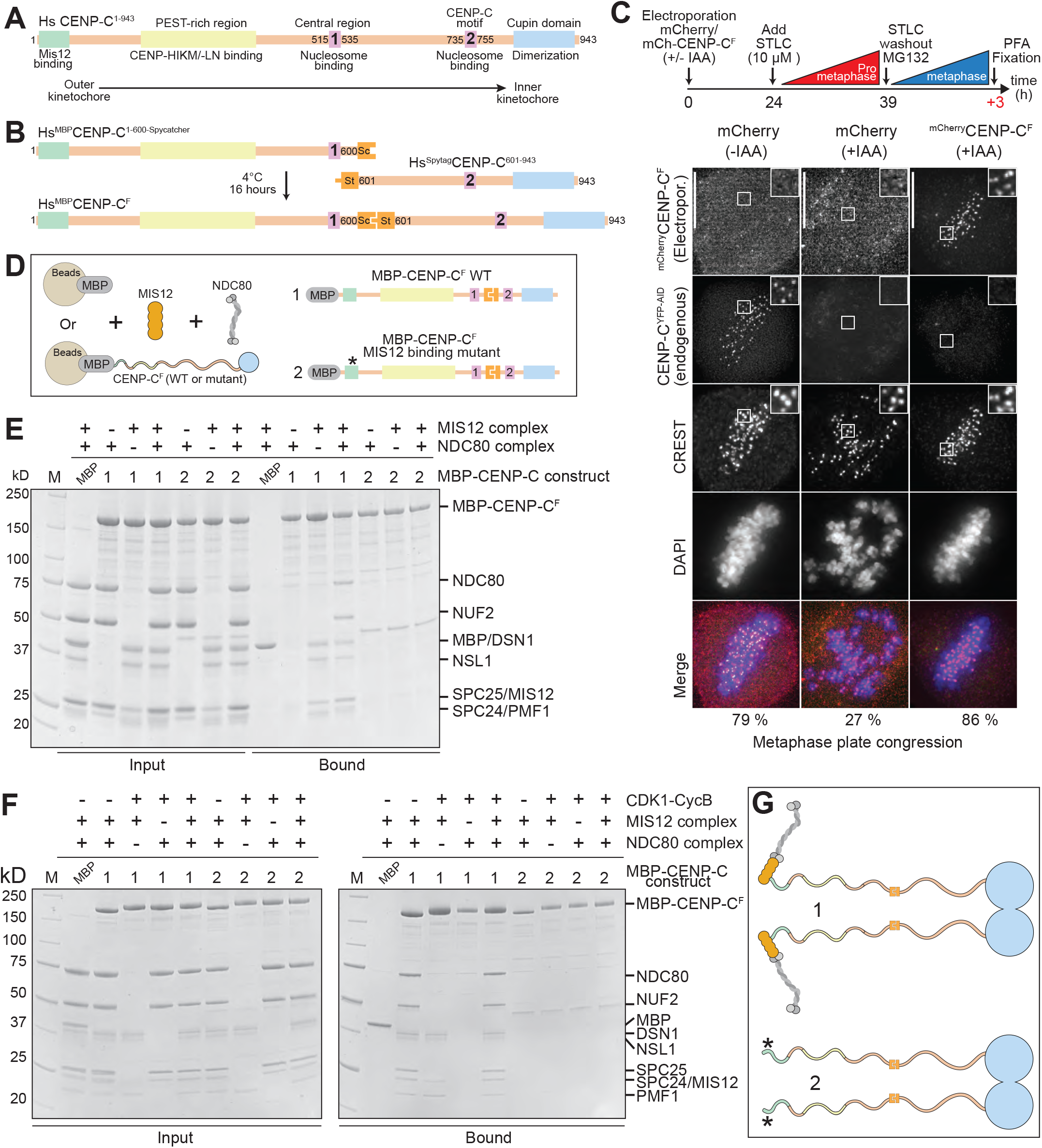
Construction and validation of a CENP-C fusion protein. **A**) Organization of human CENP-C. The N-terminal MIS12 binding region is highlighted in green, the CENP-HIKM/-LN binding region is highlighted in yellow. The central and C-terminal CENP-C motifs are highlighted in magenta. The C-terminal Cupin-like dimerization domain is highlighted in blue. The arrangement of the binding sites recapitulates the outer to inner kinetochore axis. **B**) Strategy to purify full-length CENP-C using the SpyCatcher-SpyTag system. The two individual CENP-C fragments, CENP-C^1-600^-SpyCatcher and SpyTag-CENP-C^601-943^, were incubated together to form the full-length CENP-C ligation product. **C**) Representative images showing fluorescence of electroporated mCherry or mCherry-CENP-C^F^ and endogenous CENP-C^YFP-AID^. Centromeres were visualized by CREST sera and DNA was stained by DAPI. Scale bars indicate 10 μm. Experimental regime: 24 h after electroporation cells were treated with 10 μM STLC for 15 h. STLC was washed out three times and 10 μM MG132 was added for 3 h before PFA fixation. **D**) Schematic of the performed amylose-resin pull-down assays. MBP and two MBP-CENP^F^ variants, WT (1) and MIS12 binding mutant (2), were immobilized on amylose resin as bait. MIS12C and NDC80C were added as preys. **E-F**) Result of the amylose-resin pull-down experiment. The MBP-CENP-C^F^ variant used as bait is indicated above each lane. MIS12C and NDC80C was added as indicated above each lane. MBP-CENP-C^F^ was additionally phosphorylated by CDK1-Cyclin B as indicated above each lane in F. **G**) Graphical summary of the results shown in E and F.

The functional properties of the complementary segment, CENP-C^545-943^, remain more enigmatic. The C-terminal Cupin domain (Dunwell et al., 2001) promotes dimerization of CENP-C (Chik et al., 2019; Cohen et al., 2008; Sugimoto et al., 1997; Trazzi et al., 2009; Xiao et al., 2017). The other functionally relevant segment is the CENP-C motif (indicated as ‘2’ in Figure 1A and also known as ‘conserved motif’), whose sequence is strongly related to that of the central region (Figure S1A). The CENP-C motif also binds CENP-A^NCPs^, but less robustly than the central region, and less selectively, as it binds CENP-A^NCPs^ and H3^NCPs^ with similar binding affinities (Ali-Ahmad et al., 2019; Kato et al., 2013; Watanabe et al., 2019). The significance of the duplication of nucleosome-binding motifs in HsCENP-C remains unclear, but suggests that a CENP-C dimer may interact with more than one nucleosome.

To dissect this aspect of CENP-C organization and recapitulate *in vitro* expected increases in complex stability arising from dimerization, full-length CENP-C would be invaluable, but obtaining it has proven difficult due to insolubility upon recombinant expression in different hosts (our unpublished observations). To overcome this technical hurdle, we sought to obtain full-length CENP-C from fusing two complementary segments of CENP-C with the Spy-tag/Spy-catcher system, which allows the covalent fusion of two polypeptide chains through formation of an isopeptide bond (Zakeri et al., 2012) (Figure 1B). We tested the efficiency of the fusion reaction with two complementary constructs collectively encompassing the entire CENP-C sequence, CENP-C^1-600^ and CENP-C^601-943^. These constructs respectively include the central region and the CENP-C motif, and connect through a poorly conserved fragment, thus reducing the likelihood of functional inactivation. In isolation, CENP-C^1-600^ and CENP-C^601-943^ bound with CENP-A^NCPs^ in size-exclusion chromatography (SEC) experiments (Figure S1B-C). We purified ^MBP^CENP-C^1-600-Spycatcher^ (where MBP stands for maltose-binding protein, an affinity tag; Figure S2A) and ^Spytag^CENP-C^601-943^, mixed them, and incubated them together for 16 hours to obtain the ligation product ^MBP^CENP-C^1-600-Spy-601-943^ (herewith usually referred to as ^MBP^CENP-C^F^, for CENP-C fusion; Figure S2B). The latter was separated from the excess of non-ligated fragments using SEC, obtaining a product of sufficient purity for further biochemical analyses (Figure S2C).

We performed SEC experiments to assess if ^MBP^CENP-C^F^ retains the ability to interact with CENP-A^NCPs^. ^MBP^CENP-C^F^ and CENP-A^NCPs^ interacted robustly and eluted in an apparently stoichiometric complex from the SEC column (when mixed at 1:2 molar ratio, the significance of which will be discussed more thoroughly below) (Figure S2D, yellow trace). Importantly, ^MBP^CENP-C^F^ bound only weakly to H3^NCPs^, as revealed by an only modest shift in the elution volume of H3^NCPs^, indicative of a low-affinity, rapidly dissociating interaction (Figure S2E, black trace). Thus, ^MBP^CENP-C^F^ binds selectively to CENP-A^NCPs^ over H3^NCPs^, a conclusion further supported by competition experiments with electrophoretic mobility shift assays (KW and AM, unpublished results).

### CENP-C^F^ is functional

As a stringent test to establish if CENP-C^F^ is functional, we asked if it localized to kinetochores and if it complemented the loss of endogenous CENP-C. For this, we used a previously reported colorectal adenocarcinoma DLD-1 cell line in which both endogenous CENP-C alleles were C-terminally tagged with an auxin inducible degron (AID) and EYFP (Fachinetti et al., 2015; Guo et al., 2017; Holland et al., 2012; Nishimura et al., 2009). Upon addition of the auxin derivative indole acetic acid (IAA), endogenous CENP-C was efficiently depleted within 30 to 60 minutes (Figure S3A-B), as described (Fachinetti et al., 2015; Guo et al., 2017).

Using the same fusion strategy that allowed us to obtain ^MBP^CENP-C^F^, we generated ^mCherry^CENP-C^F^, and demonstrated that it also interacts with CENP-A^NCPs^ and that it localizes to kinetochores of HeLa cells (Figure S2F-G and Figure S3C). Next, we asked if ^mCherry^CENP-C^F^ localized at kinetochores of DLD-1 cells depleted of the endogenous CENP-C^AID-EYFP^. We electroporated ^mCherry^CENP-C^F^ into DLD-1 cells undergoing a 12 h IAA treatment to achieve complete removal of the endogenous CENP-C^AID-EYFP^. ^mCherry^CENP-C^F^, but not an mCherry control, was readily identified at kinetochores, as indicated by overlapping mCherry and CREST (a centromere marker) signals, despite undetectable endogenous CENP-C^AID-EYFP^ (Figure S3D). Thus, centromere localization of ^mCherry^CENP-C^F^ did not require endogenous CENP-C. After incorporation into chromatin, CENP-A remains stably associated with centromeres even after removal of CENP-C (Cao et al., 2018; Carroll et al., 2010; Watanabe et al., 2019), suggesting that CENP-A directs re-recruitment of ^mCherry^CENP-C^F^ after electroporation in CENP-C-depleted cells.

Finally, we assessed if ^mCherry^CENP-C^F^ complemented the deleterious consequences on chromosome alignment caused by CENP-C^AID-EYFP^ depletion. 79 % of untreated cells reached metaphase alignment, against 20 % in cells treated with IAA to deplete CENP-C^AID-EYFP^ (Figure 1C). Electroporated ^mCherry^CENP-C^F^ entirely rescued the effects of CENP-C^AID-EYFP^ depletion, with >80% of ^mCherry^CENP-C^F^-positive cells displaying metaphase alignment. Collectively, these results provide strong evidence that ^mCherry^CENP-C^F^ is functional.

### CENP-C^F^ binds the outer kinetochore

Mif2p, the *S. cerevisiae* orthologue of CENP-C, adopts an auto-inhibited conformation that suppresses the ability of its N-terminal region to interact with the Mtw1C^Mis12C^ complex before binding to Cse4^CENP-A^ nucleosomes (Killinger et al., 2020). To assess if this was also true of human CENP-C^F^, we immobilized ^MBP^CENP-C^F^ and incubated it with MIS12C, NDC80C, or their combination. MIS12C bound in apparently stoichiometric amounts to CENP-C and attracted NDC80C when the complexes were combined (Figure 1D-E), arguing that the MIS12C binding motif of ^MBP^CENP-C^F^ is fully exposed. As explained below, the interaction of CENP-T with the MIS12C and NDC80C is regulated by cyclin-dependent kinase 1 (CDK1) phosphorylation. To assess if phosphorylation created additional binding sites for the outer kinetochore also on ^MBP^CENP-C^F^, we phosphorylated it with CDK1:Cyclin B. This caused a prominent electrophoretic migration shift of ^MBP^CENP-C^F^, indicative of successful phosphorylation, but did not modify the stoichiometry of outer kinetochore subunits (Figure 1F). Importantly, CENP-C binding of outer kinetochore subunits was limited to the previously identified N-terminal MIS12C binding motif (Screpanti et al., 2011), because it was ablated when the motif was mutated (construct 2, Figure 1F-G). Thus, collectively, CENP-C^F^ is not auto-inhibited and is a target of CDK1 phosphorylation, but the latter does not generate additional binding sites for outer kinetochore subunits.

### Role of CENP-C dimerization

The C-terminal Cupin domains of the *D. melanogaster* and *S. cerevisiae* CENP-Cs (DmCENP-C^1127-1410^ and Mif2^321-529^, respectively, depicted in Figure 2A) form tight dimers (Chik et al., 2019; Cohen et al., 2008; Medina-Pritchard et al., 2020; Trazzi et al., 2009). Sedimentation velocity analytical centrifugation (AUC) and SEC demonstrated that the human CENP-C Cupin domain also forms monodisperse, tight dimers (Figure 2B and Figure S4A; note that MBP is a monomer, as shown previously (Pan et al., 2017)). When expressed in interphase HeLa cells depleted of endogenous CENP-C (Figure S4B), GFP-CENP-C^721-943^ or GFP-CENP-C^760-943^ failed to localize to kinetochores, whereas weak localization was observed in presence of endogenous CENP-C (Figure S4C-F), likely because of dimerization of the cupin domain. When transfected in human cells together with full-length ^GFP^CENP-C, only Hs^mCherry^CENP-C^773-943^, but not Dm^mCherry^CENP-C^1127-1410^ nor Sc^mCherry^Mif2p^321-529^, formed dimers with Hs^GFP^CENP-C (Figure 2C) and localized, albeit weakly, to kinetochores (Figure 2D-E). Thus, Hs^GFP^CENP-C does not appreciably bind the fly or yeast CENP-C Cupin domains, revealing a strong preference for intra-species homodimerization of the Cupin domain. When expressed ectopically in HeLa cells, a construct lacking the Cupin domain, HsCENP-C^1-772^, localized very weakly to kinetochores, (Figure 2F-H and Figure S4G). Conversely, chimeric constructs of HsCENP-C^1-772^ and the fly or yeast Cupin domains, or even with the LacI dimerization domain, restored very robust kinetochore localization. We conclude that dimerization through the Cupin domain stabilizes kinetochore CENP-C, likely through enforcement of multivalent interactions with other CCAN subunits.

**Figure 2.**
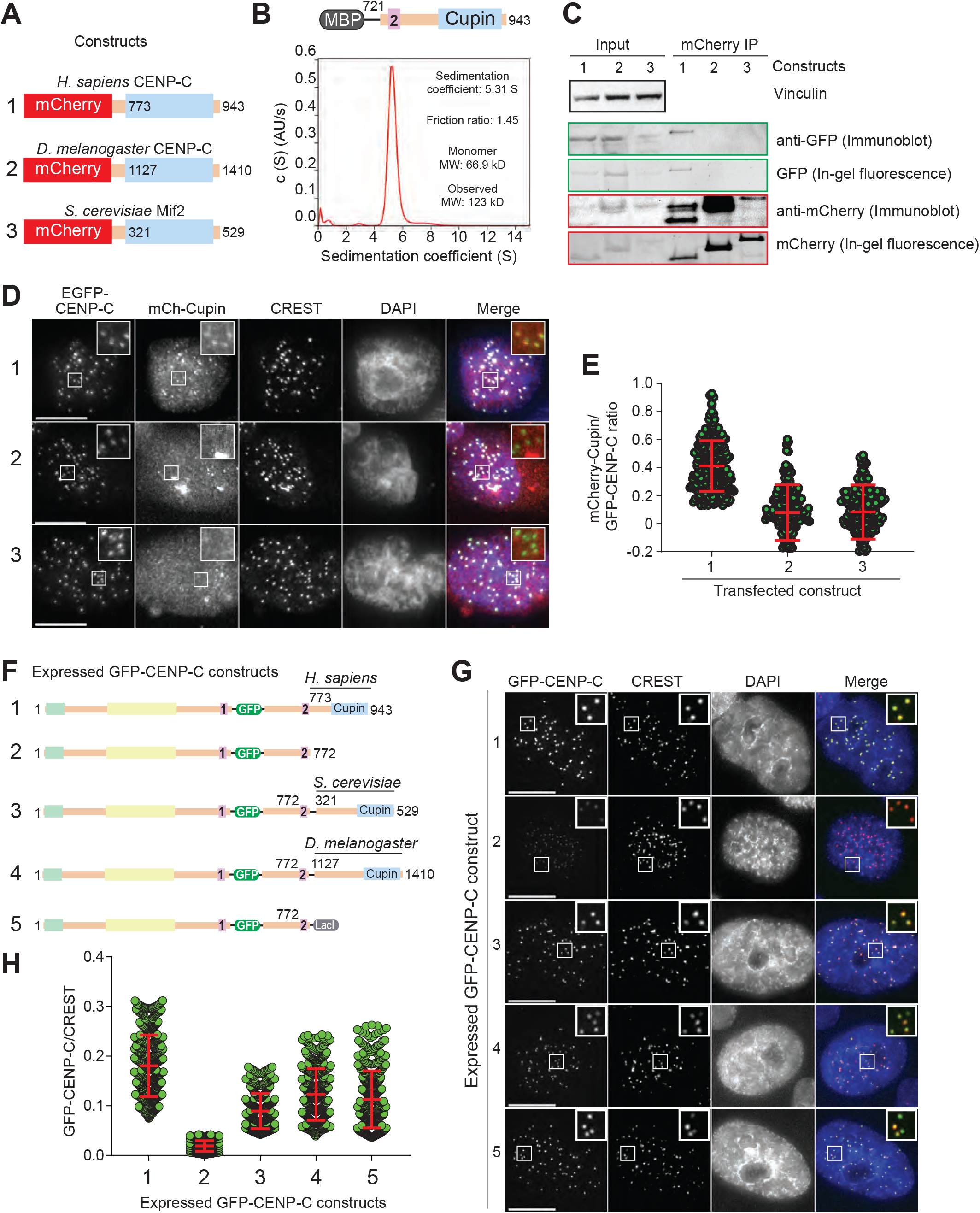
CENP-C dimerization promotes kinetochore localization efficiency. **A**) Schematic showing the three mCherry-tagged Cupin constructs from *H. sapiens*, *S. cerevisiae* and *D. melanogaster* which were co-expressed with human GFP-CENP-C in HeLa cells. **B**) Sedimentation coefficient distribution obtained from the sedimentation velocity AUC experiment using MBP-CENP-C^721-943^. The observed molecular weight of 123 kD indicates, that the sample forms a dimer. **C**) Western Blot showing the results of co-immunoprecipitation experiments using RFP-Trap beads. **D**) Representative images showing the fluorescence of GFP-CENP-C and three different mCherry-Cupin fragments as shown in A. Centromeres were visualized by CREST sera and DNA was stained by DAPI. Scale bars indicate 10 μm. **E**) Quantification of the fluorescence intensities showing the mCherry/GFP ratio. Centromeres were detected in the CREST channel using a script for semiautomated quantification. **F**) Schematic showing the different GFP-CENP-C variants which were expressed in HeLa cells. **G**) Representative images showing the fluorescence of the different GFP-CENP-C variants as shown in F. Centromeres were visualized by CREST sera and DNA was stained by DAPI. Scale bars indicate 10 μm. **H**) Quantification of the GFP fluorescence intensities.

### CENP-C^F^ binds two CENP-A nucleosomes

Because CENP-C^F^ encompasses two nucleosome-binding motifs, its dimerization through the Cupin domain generates a dimer with four nucleosome-binding motifs. Due to its pseudo-two-fold symmetric organization, a nucleosome core particle (NCP) may contain two binding sites for the CENP-C motifs, so that a CENP-C homodimer might be expected to interact with at least two NCPs, or more if further oligomerization came into play. To test this, we performed SEC experiments to assess how CENP-A^NCPs^ interacts with CENP-C^F^. As controls, we also included three CENP-C^F^ variants in which the central region, the CENP-C motif, or both had been mutated to impair CENP-A^NCP^ binding (Kato et al., 2013). Point mutations directed against conserved residues in the central region or in the CENP-C motif (CENP-C^1-600-R522A/W531A^ or CENP-C^601-943-R742A/W751A^, respectively) abolished CENP-A binding (Figure S1B-C), indicating that each set of mutations is highly penetrant in isolation.

CENP-A^NCPs^ combined with ^MBP^CENP-C^F^ at 2:1 molar ratio (5 μM CENP-A^NCP^ and 2.5 μM CENP-C, dimer concentration) eluted from the SEC column almost entirely in one peak, with no residual free CENP-A^NCPs^ (Figure 3A,E and Figure S5A), implying that the four CENP-C motifs in the dimeric CENP-C^F^ provide binding sites for at least two CENP-A^NCPs^. Conversely, CENP-C containing two non-functional CENP-C motifs did not interact with CENP-A at all (Figure 3B,E and Figure S5B). Importantly, CENP-C^F-R522A/W531A^ and CENP-C^F-R742A/W751A^ bound only one equivalent of CENP-A^NCP^, rather than two as CENP-C^F^ (Figure 3C-D,E and Figure S5C-D). These stoichiometries were inferred both from the absorbance intensities of the peaks eluting from the SEC column, and from the intensity of histone bands in Coomassie-stained SDS-PAGE of the CENP-C^F^-bound fractions (Figure 3E).

**Figure 3.**
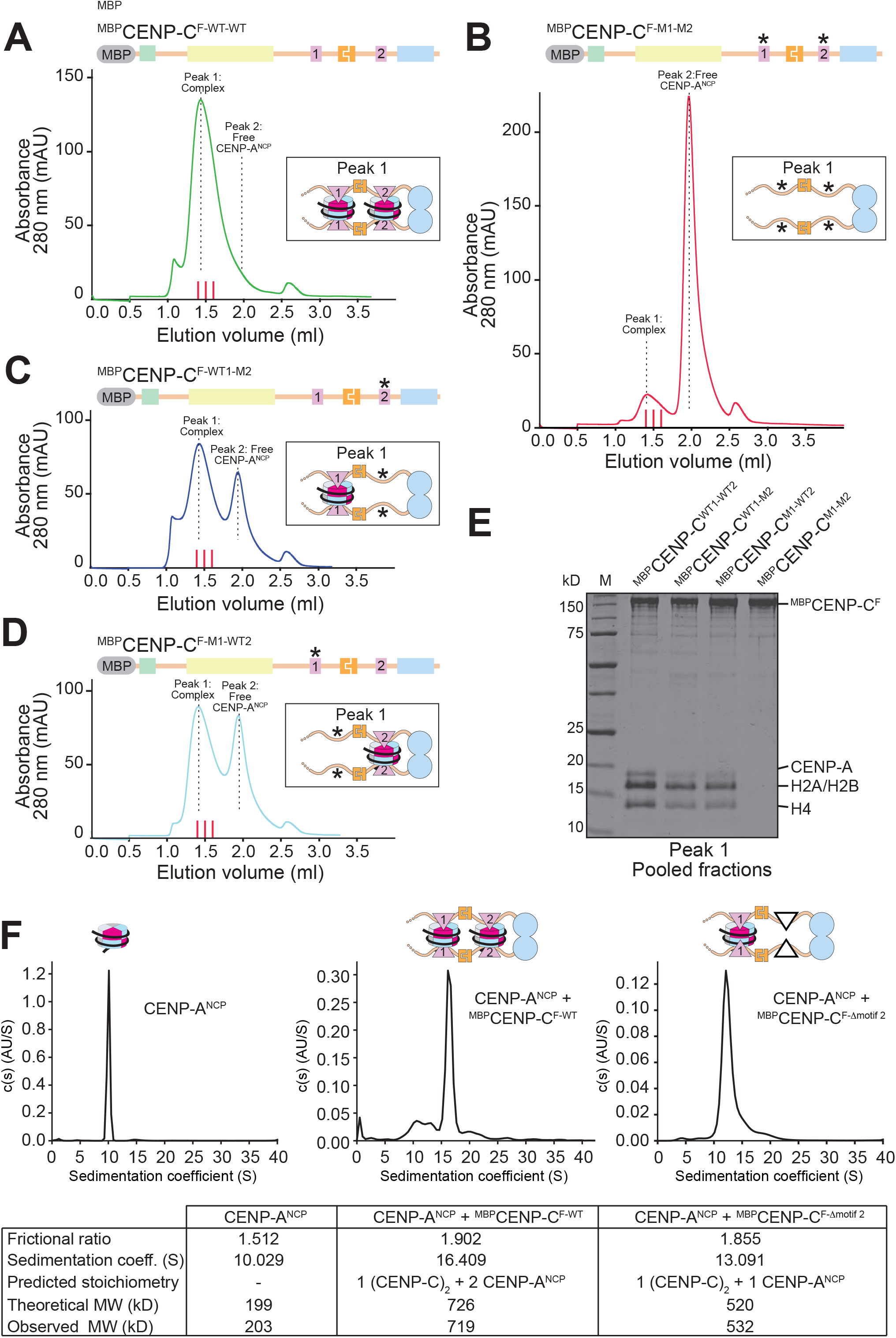
CENP-C binds two CENP-A nucleosomes. **A-D**) Analytical SEC experiments were performed to demonstrate the interactions between different CENP-C variants and CENP-A^NCP^. Red lines between 1.4 and 1.6 ml elution volume indicate fractions collected for SDS-PAGE analysis shown in E. The SEC chromatograms and SDS-PAGEs of all individual components are shown in Figure S5. **E**) SDS-PAGE result showing the pooled fractions of the peak 1 complexes shown in the SEC chromatograms in A-D. **F**) Sedimentation coefficient distributions obtained from the sedimentation velocity AUC experiments using CENP-A NCP (left), CENP-A NCP + MBP-CENP-C^F-WT^ (middle) and CENP-A NCP + CENP-C^F-Dmotif2^ (right). The table summarizes the results of the three experiments.

In an orthogonal approach, we used sedimentation velocity analytical ultracentrifugation (AUC) to assess the molecular mass of the CENP-C^F^:CENP-A^NCP^ complex obtained at the 2:1 molar ratio. These experiments revealed an excellent agreement between the expected and predicted molecular mass for a complex containing a CENP-C^F^ dimer and two CENP-A^NCPs^, with apparently little additional contamination from complexes of different stoichiometry (Figure 3F and Figure S6A-D). Furthermore, deletion of the CENP-C motif resulted in a complex containing a single CENP-A^NCP^, again with an excellent agreement between the theoretical and expected molecular masses. Collectively, these experiments indicate that a CENP-C^F^ dimer has an intrinsic potential to bind two CENP-A^NCPs^, without forming significant amounts of additional higher molecular weight species that might potentially arise when combining multivalent binding partners like CENP-C and NCPs.

### CENP-C interacts with CENP-A also through the CENP-HIKMLN complex

CENP-N, like CENP-C, interacts directly with CENP-A^NCPs^ by recognizing the CENP-A L1 loop (Allu et al., 2019; Cao et al., 2018; Chittori et al., 2018; Pentakota et al., 2017; Tian et al., 2018). CENP-C, on the other hand, interacts with the acidic patch of H2A-H2B and with the CENP-A C-terminal tail (Ali-Ahmad et al., 2019; Allu et al., 2019; Cao et al., 2018; Carroll et al., 2010; Kato et al., 2013; Weir et al., 2016; Yan et al., 2019). These features may allow CENP-C and CENP-N to bind concomitantly to their cognate binding sites on the same CENP-A nucleosome (Allu et al., 2019), although an alternative binding model where CENP-N occupies a non-canonical position has been recently proposed (Yan et al., 2019). Because CENP-NL (and CENP-HIKM) interact tightly with CENP-C also through the PEST region (Klare et al., 2015; Pentakota et al., 2017) (Figure 1A), CENP-C may be expected to enable an additional contact of CCAN with the CENP-A nucleosome through CENP-N.

To investigate this possibility, we immobilized three MBP-tagged CENP-C^F^ variants on amylose beads and used them as baits with CENP-A^NCPs^ and CCAN sub-complexes (Figure 4A). Specifically, we used CENP-C^F^, CENP-C^F-R522A/W531A/R742A/W751A^ (indicated as ‘nucleosome binding mutant’), and a nucleosome binding mutant additionally impaired in the interaction with CENP-NL and CENP-HIKM through previously described point mutations (Klare et al., 2015; Pentakota et al., 2017) (and indicated as ‘nucleosome and CCAN binding mutant’). We then incubated these baits with saturating concentrations of CENP-A^NCPs^, CENP-HIKMLN, or their combination. After washing, species retained on beads were resolved by SDS-PAGE and visualized by Coomassie staining. This revealed that CENP-C^F^ binds the same levels of CENP-A^NCPs^ and CENP-HIKMLN when presented individually or concomitantly (Figure 4A). Conversely, the CENP-A binding mutant of CENP-C^F^ did not bind CENP-A^NCPs^, but retained binding to CENP-HIKMLN. When CENP-A^NCPs^ and CENP-HIKMLN were co-incubated with this mutant, binding of CENP-A^NCPs^ was restored, but at reduced levels. The residual CENP-A^NCP^ was binding to this mutant through CENP-N in the CCAN complex, because the CENP-A and CCAN mutant of CENP-C^F^ bound only residual levels of CENP-A^NCPs^ or CENP-HIKMLN (Figure 4A, quantified in Figure S6E). Thus, binding of CENP-C^F^ to other CCAN subunits allows a robust point of contact with the CENP-A nucleosome through CENP-N (and possibly additional still undiscovered points of contact within the CENP-HIKMLN complex) even in the absence of functional CENP-A binding motifs.

**Figure 4.**
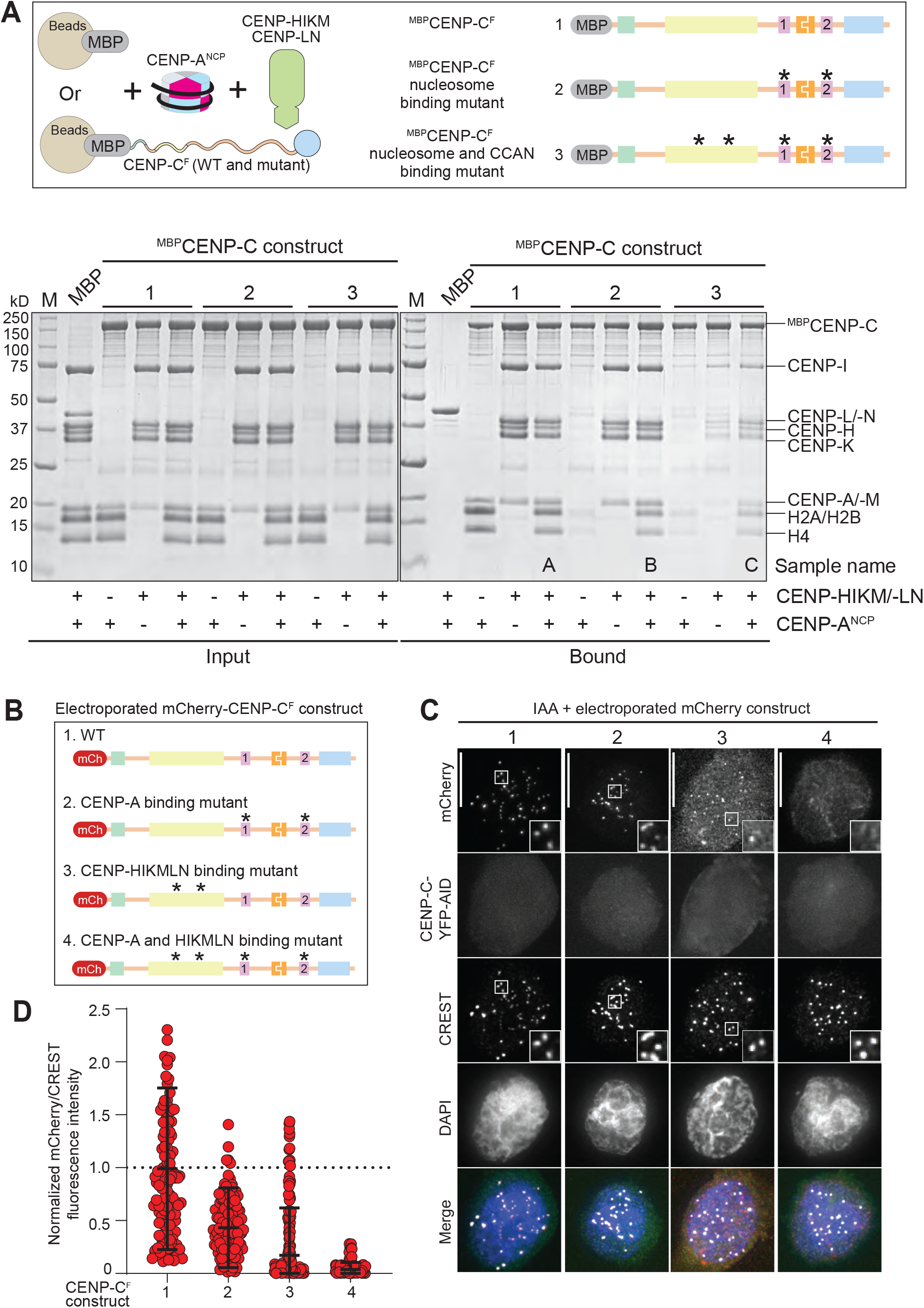
Relative contributions of CENP-C and CCAN to CENP-A binding. **A**) Schematic of the performed amylose-resin pull-down assays. MBP and three MBP-CENP^F^ variants, WT (1), Nucleosome binding mutant (2) and Nucleosome and CCAN binding mutant (3), were immobilized on amylose resin as bait. CENP-A NCP and the CCAN members CENP-HIKM/-LN were added as prey. The two SDS-PAGEs show the results of the amylose-resin pull-down experiment. The MBP-CENP-C^F^ variant used as bait is indicated above each lane. CENP-A NCP and CENP-HIKM/-LN was added as indicated below each lane. The left gel shows the input fractions, the right gel shows the bound fractions. **B**) Schematic showing the different electroporated mCherry-CENP-C^F^ variants. **C**) Representative images showing the fluorescence of endogenous CENP-C-YFP-AID and the electroporated mCherry-CENP-C^F^ variants as shown in B. Centromeres were visualized by CREST sera and DNA was stained by DAPI. Scale bars indicate 10 μm. **D**) Quantification of the mCherry fluorescence intensities normalized to the centromere marker CREST and to mCherry-CENP-C^F-WT^.

Kinetochore localization of electroporated ^mCherry^CENP-C^F^ in cells depleted of endogenous CENP-C^YFP-AID^ in DLD-1 cells demonstrated the main predictions from the *in vitro* assay (Figure 4B-D). Specifically, CENP-C^F^ localized robustly to kinetochores, whereas the CENP-A binding mutant showed a partial localization impairment. Likely, this mutant is targeted to the centromere by CENP-HIKM or CENP-LN, because the CCAN binding mutant, already in isolation, showed very strongly impairment of kinetochore localization. This was compounded by additional mutation of the CENP-A binding motifs, which made the construct undetectable at the kinetochore. Collectively, these observations indicate that the interaction of CENP-C with centromeres is cooperative and reflects its multiple binding sites for CENP-A and other CCAN subunits.

### The interaction of CENP-T with CCAN

The CCAN subunits CENP-S, CENP-T, CENP-W, and CENP-X are histone-fold domain (HFD) proteins that associate in stable dimeric pairs (CENP-SX and CENP-TW) and as tetramer (Hori et al., 2008). We reconstituted the CENP-TWSX complex to identify conditions for its stable incorporation in reconstituted kinetochores. In solid phase binding assays with immobilized ^MBP^CENP-C^F^ (Figure 5A-B) we observed binding of CENP-TWSX in presence of CENP-HIKM and CENP-LN, reduced binding upon omission of CENP-LN, and poor binding upon omission of CENP-HIKM or of both CENP-HIKM and CENP-LN (shown schematically in Figure 5C). Thus, CENP-HIKM, and marginally also CENP-LN, provide a binding platform for CENP-TWSW, in agreement with previous studies in *S. cerevisiae* (Basilico et al., 2014; Hinshaw and Harrison, 2020; Pekgoz Altunkaya et al., 2016; Zhang et al., 2020). A highly conserved C-terminal helical hairpin of CENP-T (helices α4 and α5), which extends the HFD to complete a binding interface for the CENP-HIK complex (Hinshaw and Harrison, 2020; Zhang et al., 2020), was required for tight binding to the ^MBP^CENP-C^F^-CCAN complex *in vitro* as well as for its kinetochore recruitment after electroporation (Figure 5D-E and Figure S6F-G).

**Figure 5.**
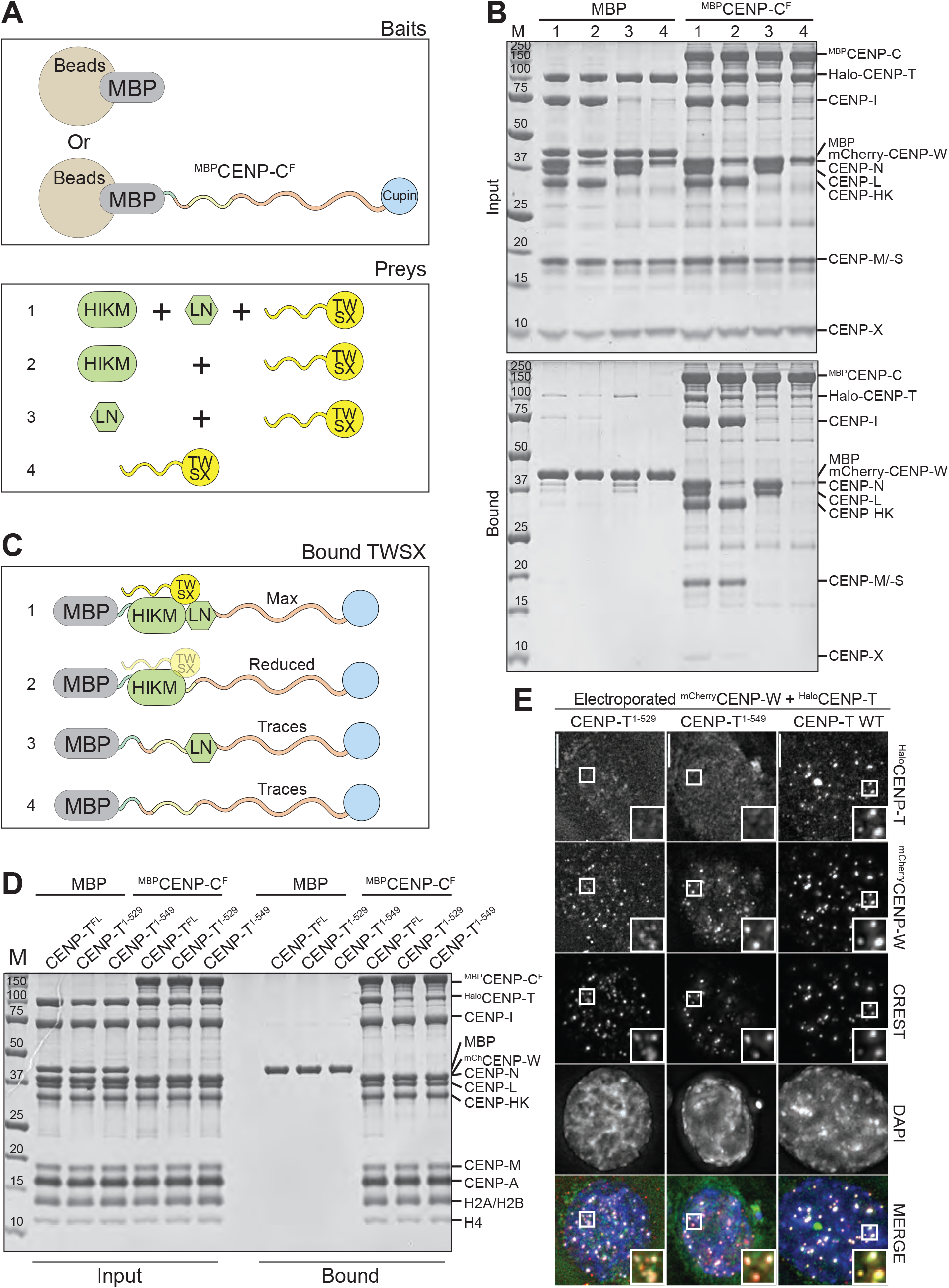
Determinants of CENP-T binding to CCAN. **A**) Schematic of the performed amylose-resin pull-down assays. MBP and MBP-CENP^F^ were immobilized on amylose resin as bait. CENP-TWSX was added as prey, either alone or in combination with CENP-HIKM and/or CENP-LN. **B**) Result of the amylose-resin pull-down experiment. The used preys are indicated above each lane. The top gel shows the input fractions, the bottom gel shows the bound fractions. **C**) Graphical summary of the results obtained in B. **D**) SDS-PAGE result showing an amylose-resin pull-down experiment, in which three different Halo-tagged CENP-T variants were used, namely CENP-T^1-529^, CENP-T^1-549^ and CENP-T^FL^. MBP-CENP-C was used as bait and MBP was used as negative control. **E**) Representative images showing the fluorescence of mCherry-CENP-W and three Halo-tagged CENP-T variants labelled with a green fluorescent ligand. Centromeres were visualized by CREST sera and DNA was stained by DAPI. Scale bars indicate 10 μm.

### Role of CENP-T in outer kinetochore assembly

CENP-T, like CENP-C, is crucial for outer kinetochore assembly (Gascoigne et al., 2011; Hara et al., 2018; Huis In ’t Veld et al., 2016; Lang et al., 2018; Malvezzi et al., 2013; Nishino et al., 2013; Pekgoz Altunkaya et al., 2016; Schleiffer et al., 2012; Suzuki et al., 2015). The flexible N-terminal region of human CENP-T binds two distinct NDC80 complexes through motifs encompassing the CDK1 targets Thr11 and Thr85 (Huis In ’t Veld et al., 2016). A third CDK1 phosphorylation site, Ser201, possibly with an additional contribution from Thr195, is further responsible for an interaction with an entire KMN network complex through a binding site on MIS12C (Huis In ’t Veld et al., 2016; Rago et al., 2015). Thus, CENP-T recruits two NDC80Cs directly and one KMN (which contains an additional NDC80 complex) while CENP-C, as shown above, recruits a single KMN network through MIS12C.

CENP-C binds MIS12C at the interface between the Head1 region (encompassing the N-terminal regions of the MIS12 and PMF1 subunits) and a “connector” encompassing the α3 helices of NSL1 and DSN1 (Petrovic et al., 2016). The regions of NSL1 and DSN1 preceding the connector form a globular domain named Head2 (Figure 6A-B). A disordered segment of DSN1 preceding Head2 contains at least two Aurora B target sites (Ser100^DSN1^ and Ser109^DSN1^) that control access of CENP-C through an intramolecular regulatory mechanism (Akiyoshi et al., 2013; Bonner et al., 2019; Dimitrova et al., 2016; Kim and Yu, 2015; Petrovic et al., 2016; Rago et al., 2015; Welburn et al., 2010). CENP-C binds MIS12C with ~150-fold higher affinity after Aurora B-mediated phosphorylation of Ser100^DSN1^ and Ser109^DSN1^ (Petrovic et al., 2016). Deletions of the entire N-terminal region of DSN1 and NSL1 including Head2 (indicated simply as MIS12C^ΔHead2^) or of a 10-residue segment encompassing the two phosphorylation sites (MIS12C^Δ10^) bypass the requirement for phosphorylation and allow unhindered access to CENP-C (Kim and Yu, 2015; Petrovic et al., 2016; Rago et al., 2015). Previously, we showed that CENP-T and CENP-C cannot be contemporarily bound to the same MIS12C (Huis In ’t Veld et al., 2016), suggesting partial or complete overlap of their binding sites on MIS12C. Whether access of CENP-T to MIS12C is also regulated by Aurora B activity, however, is unknown. More generally, due to lack of structural information on the CENP-T:MIS12C complex, as well as of obvious resemblances in the MIS12C-binding regions of CENP-C and CENP-T, how CENP-T binds MIS12C remains unclear.

**Figure 6.**
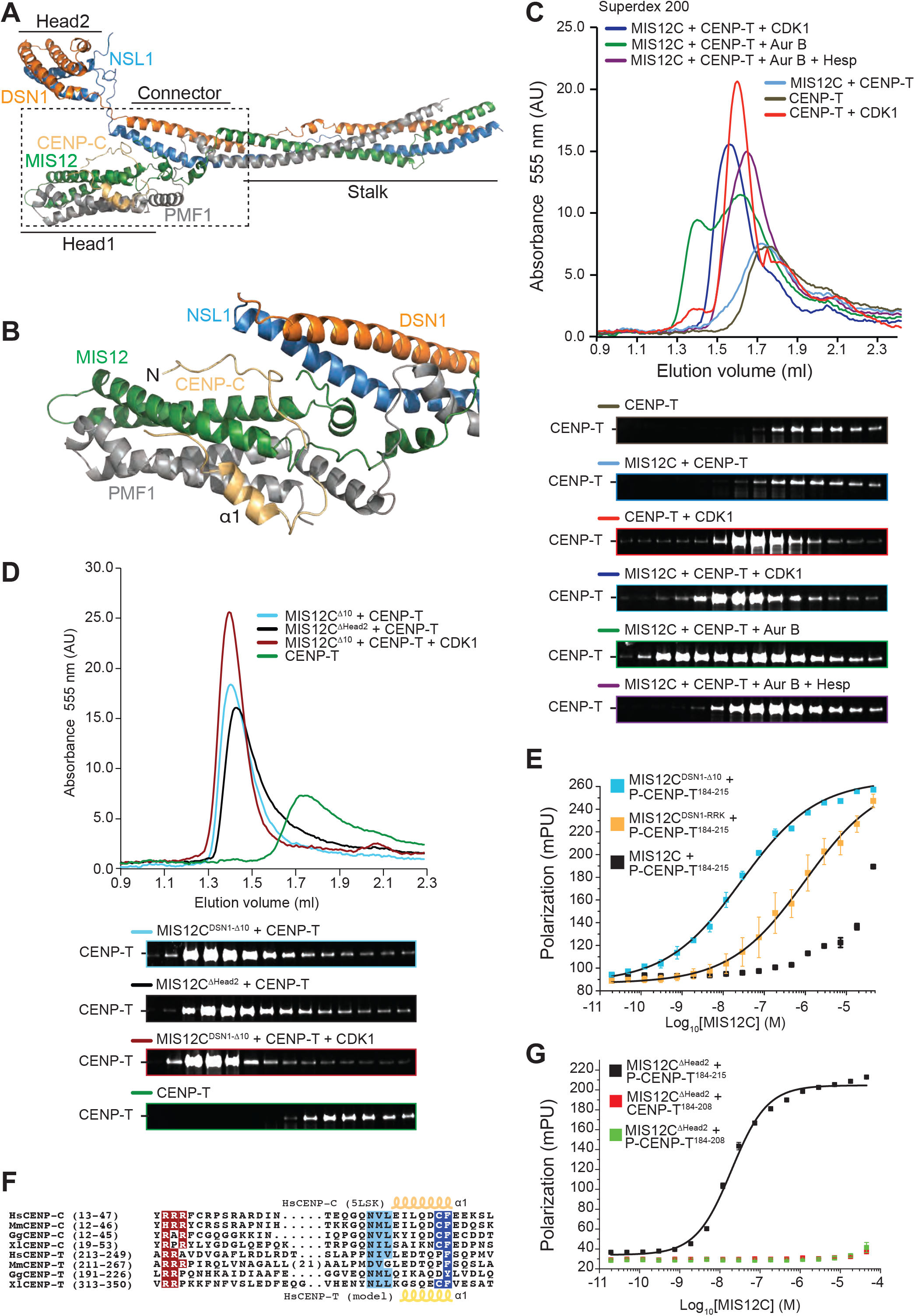
Molecular mechanism of the CENP-T:MIS12 interaction. **A**) Cartoon model of the human MIS12 complex in complex with the CENP-C N-terminal region (PDB ID 5LSK). **B**) Close-up of the MIS12:CENP-C interaction interface. **C**) Size-exclusion chromatography on a Superdex 200 column of the indicated species. **D**) Size-exclusion chromatography on a Superdex 200 column of the indicated species. **E**) Binding isotherms by fluorescence polarization with a fluorescent Ser201-phosphorylated CENP-T^184-215^ peptide and the indicated MIS12 complex variants. **F**) Multiple sequence alignment of segments of the MIS12 complex-binding regions of CENP-C and CENP-T from the indicated species (Hs, *Homo sapiens*; Mm, *Mus musculus*; Gg, *Gallus gallus*; Xl, *Xenopus laevis*). **G**) Binding isotherms by fluorescence polarization obtained with the indicated fluorescent S201-phosphorylated or non-phosphorylated CENP-T peptides and the MIS12^ΔHead2^ mutant complex.

To investigate this, we monitored the elution of fluorescently labelled CENP-T^1-373^ (Huis In ’t Veld et al., 2016) from a SEC column upon addition of MIS12C. No binding of CENP-T^1-373^ to MIS12C was observed, as the elution profile of fluorescent CENP-T^1-373^ was essentially identical to that of CENP-T^1-373^ in isolation (Figure 6C). In absence of MIS12C, CDK1 phosphorylation caused CENP-T^1-373^ to elute at an earlier volume, possibly because phosphorylation causes decompaction of the N-terminal domain. CDK1 phosphorylation also enhanced CENP-T^1-373^ binding to MIS12C, as shown by a further decrease of the CENP-T^1-373^ elution volume. A more significant shift caused by addition of Aurora B was abrogated in the presence of the Aurora B inhibitor Hesperadin (Figure 6C). MIS12C^DSN1-Δ10^ or MIS12C^ΔHead2^, which lack the Aurora B phosphorylation motif, caused a more pronounced shift in the CENP-T^1-373^ elution volume, that was further reinforced after addition of CDK1 (Figure 6D). Thus, phosphorylation of CENP-T and MIS12C by CDK1 and Aurora B, respectively, or deletion of the Aurora B target motif on MIS12C, increase the binding affinity of the MIS12C:CENP-T complex, as previously observed with CENP-C (Kim and Yu, 2015; Petrovic et al., 2016; Rago et al., 2015). Accordingly, a fluorescent synthetic peptide encompassing residues 184-215 of CENP-T and phosphorylated on Ser201 (P-CENP-T^184-215^) bound to MIS12C^DSN1-Δ10^ with a dissociation constant of ~27 nM in fluorescence anisotropy binding affinity measurements, approximately 1000-fold better than to control MIS12C (Figure 6E and Table S1).

Arginine residues in the Aurora B target motif of DSN1 are important for the auto-inhibition mechanism, and their mutation to alanine (MIS12C^DSN1-RRKAAA^) is sufficient to enhance binding to CENP-C (Petrovic et al., 2016). We therefore speculated that these arginine residues compete directly with equivalent positively-charged residues in the N-terminal region of CENP-C (Arg14-Arg15-Arg16; Figure 6F) until Aurora B phosphorylation of DSN1 at neighboring serines neutralizes charge (Petrovic et al., 2016). Because MIS12C^DSN1-RRKAAA^ also bound to the P-CENP-T^184-215^ peptide more tightly than control MIS12C (Figure 6E), we speculated that a conserved short stretch of positive charges in CENP-T, Arg214 and Arg215 (Figure 6F) plays a role equivalent to that of Arg14-Arg16 in CENP-C. In agreement with this idea, a peptide excluding this region (CENP-T^184-208^ instead of P-CENP-T^184-215^) did not bind MIS12C^ΔHead2^, regardless of its phosphorylation status (Figure 6G). CENP-C binds MIS12C also through the α1-helix downstream from the arginine-rich motif (Figure 6B). A segment with helical propensity around residues 238-245 of CENP-T can be tentatively aligned to the CENP-C α1 helix (Figure 6F). Collectively, these results suggest that residues 213-249 of CENP-T bind MIS12C similarly to residues 14-48 of CENP-C, as modelled in Figure 7A-B.

**Figure 7.**
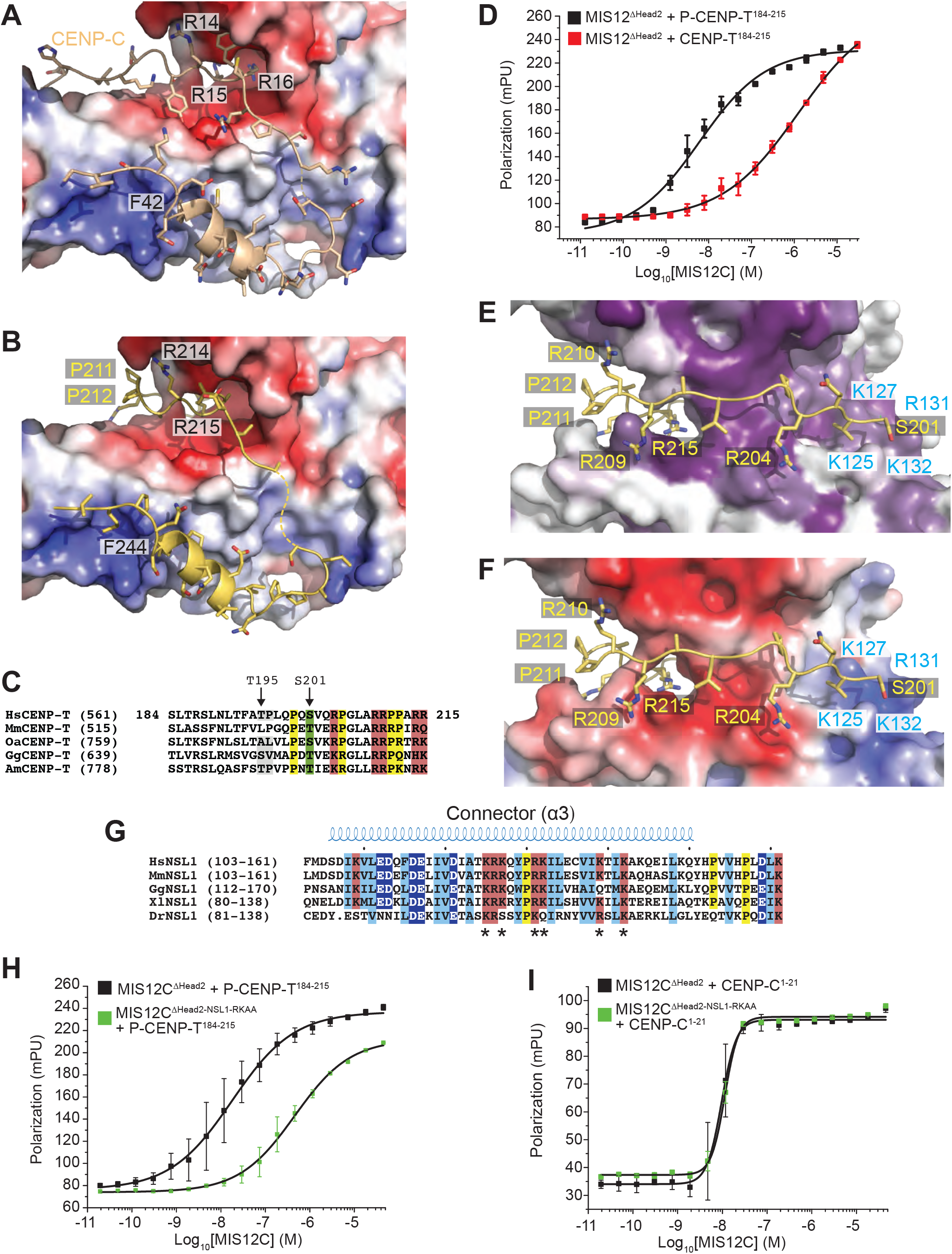
Structural basis of CENP-T:MIS12 interaction. **A**) Surface electrostatics displayed on the MIS12 complex at the interface with CENP-C (wheat). **B**) Structural model of the CENP-T peptide (residues 213-249) interaction with the MIS12C using the CENP-C N-terminal region as a template (PDB ID 5LSK) and the sequence alignment shown in Figure 6F. CENP-T was modelled on the same interface. **C**) Multiple sequence alignment of CENP-T in the unique region containing the Thr195 and Ser201 phosphorylation sites, and encompassing the modelled residues (201-212) shown in E and F, which precede the region aligned on CENP-C (Figure 6F). **D**) Binding isotherms by fluorescence polarization obtained with the indicated fluorescent S201-phosphorylated or non-phosphorylated CENP-T peptides and the MIS12^ΔHead2^ mutant complex. **E**) Structural model of the CENP-T^201-212^:MIS12C interaction and the MIS12C surface conservation of residues lining the possible CENP-T binding site. **F**) Charge distribution on the same interface discussed in H. **G**) Sequence analysis of the connector region of NSL1 pinpointing multiple conserved positively charged residues potentially involved in the phospho-dependent recognition of CENP-T. **H**) Binding isotherms by fluorescence polarization obtained with the indicated fluorescent S201-phosphorylated peptide and the MIS12^ΔHead2^ control complex or the MIS12^ΔHead2^ with alanine mutations at Arg131^NSL1^ and Lys132^NSL1^. **I**) As in H, but with a fluorescent CENP-C^1-21^ peptide.

Phosphorylation of the CENP-T^184-215^ peptide on Ser201 (Figure 7C) enhanced the binding affinity for MIS12C^ΔHead2^ approximately 200-fold (Figure 7D and Table S1). Chemical crosslinking coupled with mass spectrometry on the MIS12C:P-CENP-T^1-373^ complex (unpublished observations) identified crosslinks between Ser201^CENP-T^ and Lys127^NSL1^ or Lys103^MIS12^ that allowed us to restrain the orientation of the CENP-T peptide on the Mis12C surface and model the ultimate path of the peptide (residues 201 to 212, Figure 7E-F). Lys127^NSL1^ and Lys103^MIS12^ are part of a positively charged, conserved surface ridge residing at the junction between the connector and Head1, opposite to the CENP-C binding site, and also comprising Arg131^NSL1^ and Lys132^NSL1^ (Figure 7E-G). Mutation of the latter two residues to alanine (MIS12C^NSL1-RKAA^) reduced binding affinity for the P-CENP-T^184-215^ peptide approximately 25-fold (Figure 7H and Table S1), strongly implicating them in recognition of P-Ser201^CENP-T^. This model of the interaction of CENP-T with MIS12C predicts that two additional residues, Lys139^NSL1^ and Lys142^NSL1^, are well positioned to interact with P-Thr195^CENP-T^ (Huis In ’t Veld et al., 2016). In the structure of the MIS12C:CENP-C complex, this ridge is not involved in CENP-C binding (Petrovic et al., 2016). Accordingly, a CENP-C^1-21^ peptide bound MIS12C^WT^ and MIS12C^NSL1-RKAA^ with essentially identical affinity (Figure 7I). Collectively, these data suggest that the CENP-T chain occupies a unique MIS12C ridge until it bends, around Pro211 and Pro212, to interact with the previously identified CENP-C binding interface. This model of the CENP-T-MIS12C interaction predicts the observed competitive binding to MIS12C of CENP-C and CENP-T.

### Reconstitution of a complete kinetochore module

Finally, we aimed to assemble a complete kinetochore module comprising all the inner and outer kinetochore subunits. To monitor the incorporation of NDC80C in the reconstituted particles, we used wild type CENP-T (Figure 8A, construct 2) or a CENP-T mutant where the two direct NDC80C binding sites, Thr11 and Thr85, were mutated to alanine (construct 1). We then incubated these constructs as baits on solid phase with excess NDC80C and MIS12C and CDK1:Cyclin B (Figure 8B; the outcome of this experiment is presented schematically in Figure 7C and quantified in Figure 8D and Figure S7D). Without MIS12C, construct 1 bound only background levels of NDC80C (sample A). Addition of MIS12C (sample B) resulted in the binding of one equivalent of NDC80C. Construct 2, on the other hand, bound approximately two equivalents of NDC80C in the absence of MIS12C (sample C), and three equivalents when MIS12C was also added (sample D). These stoichometries suggest that phosphorylation of CENP-T on Thr11, Thr85, and Ser201 is complete and leads to saturation of these binding site.

**Figure 8.**
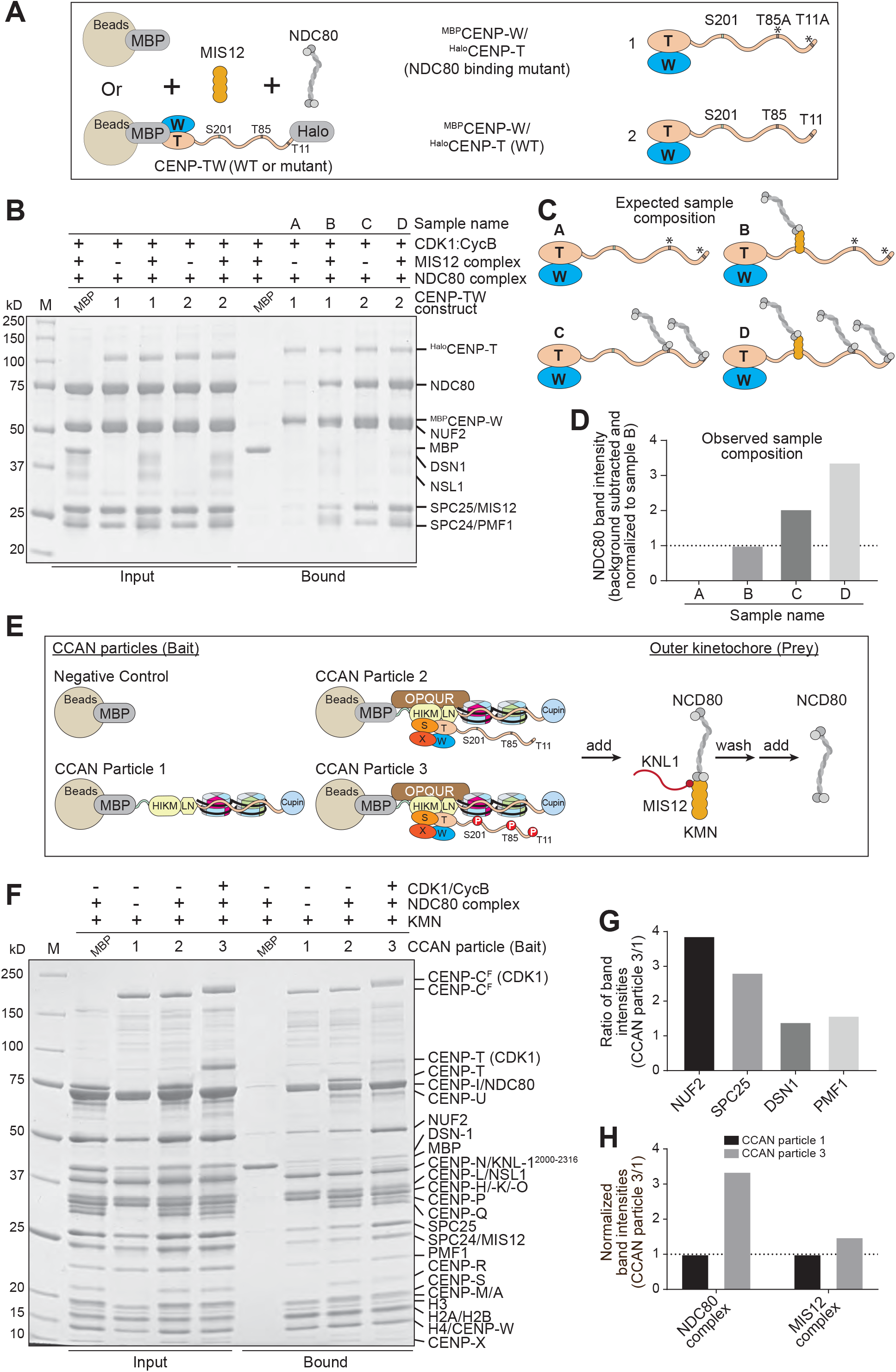
Reconstitution of a complete human kinetochore module. **A**) Schematic of the performed amylose-resin pull-down experiments. Two MBP-CENP-W/Halo-CENP-T variants, Halo-CENP-T WT (1) and Halo-CENP-T^T11AT85A^ (2) were immobilized on amylose beads as bait. NDC80C was added as prey, either alone or in combination with MIS12C. **B**) SDS-PAGE result of the performed pull-down experiment. The MBP-CENP-W/Halo-CENP-T construct used as bait is indicated above each lane. Preys were added as indicated above each lane. All samples were phosphorylated by CDK1/Cyclin B during the experiment. **C**) Graphical summary of the expected sample composition of the four samples shown in B. **D**) Quantification of the NDC80 band intensities of the four samples shown in B. The NDC80 band intensity of sample A was subtracted as background. All intensities were normalized to sample B. **E**) Schematic of the performed amylose-resin pull-down experiment. MBP and three CCAN particles assembled on MBP-CENP-C were immobilized on amylose resin as bait. The outer kinetochore KMN complex was added to the bait and incubated for 1 hour. Subsequently, unbound KMN complex was washed away and NDC80 complex was added. After a 1 hour-incubation, unbound NDC80 complex was washed away and the samples were analyzed by SDS-PAGE. **F**) SDS-PAGE result of the performed amylose-resin pull-down experiment. The CCAN particle used as bait is indicated above each lane. Samples were phosphorylated by CDK1/Cyclin B as indicated. **G**) Quantified ratio of band intensities of the indicated proteins shown in the bound fractions in F. The band intensity quantified in complex 3 was divided by the band intensity quantified in complex 1. **H**) Quantified intensities of the NDC80 complex and the MIS12 complex normalized to CCAN particle 1. The quantified NDC80 complex intensity represents the average of the NUF2 and SPC25 band intensities. The quantified MIS12 complex intensity represents the average of the DSN1 and PMF1 band intensities.

To obtain complete kinetochore particles, we reconstituted three ^MBP^CENP-C^F^ baits with tandem CENP-A^NCPs^-H3^NCPs^ dinucleosomes built on two consecutive human α-satellite repeats, and further decorated them with a different complement of CCAN subunits (particles 1 to 3 in Figure 8E; see also Figure S7A-C). Particle 3 contained all the CCAN subunits and was additionally phosphorylated by CDK1:Cyclin B. We first incubated the immobilized baits with KMN network (generated with the MIS12^Δ10^ mutant that binds maximally to CENP-C or CENP-T), and after washing the excess of unbound KMN we added an excess of free NDC80C. After further washing, we analyzed bound proteins by SDS-PAGE and Coomassie staining (Figure 8F). CDK1:Cyclin B phosphorylation was effective, because it caused large shifts in the mobility of ^MBP^CENP-C^F^ and CENP-T. We quantified band intensities of scanned gels to determine the apparent stoichiometries of outer kinetochore subunits. The intensity of KMN subunits in Particle 1, whose only linkage to the KMN is through the ^MBP^CENP-C^F^ baits, were taken as reference to assess the effects of integrating CENP-T and other CCAN subunits. We then examined the intensities of KMN subunits in particle 3 and compared them with those in particle 1. The ratio of particle 3/particle 1 MIS12C subunits (averaged over two subunits, DSN1 and PMF1), was approximately 1.5 (Figure 8G-H and Figure S7D-H), compatible with the addition of a single equivalent of CENP-T, if CENP-T remained fully saturated with MIS12C on the beads, as supported by the experiments in Figure 8B. Alternatively, the observations are compatible with the addition of two equivalents if CENP-T underwent partial dissociation from MIS12C during the experiment. On the other hand, the particle 3/particle 1 ratio of NDC80C subunits (also averaged over two subunits, NUF2 and SPC25) was higher than 3, compatible with the addition of two equivalents of CENP-T with full saturation of the Thr11 and Thr85 sites and partial saturation of the KMN binding site (Figure 8G-H and Figure S7D-H).

## Discussion

We have reconstituted a particle incorporating all known core subunits of the human kinetochore, mostly in full length form (Figure 9A). The outer kinetochore subunits assemble on this particle with stoichiometries approaching the calibrated fluorescence intensity measurements performed on entire human kinetochores (Suzuki et al., 2015). CENP-TW, however, appeared to be generally sub-stoichiometric in our reconstitutions, even in presence of nucleosomes, explaining why stoichiometries in our particles don’t yet reach the highest ratios of outer to inner kinetochore subunits observed at human kinetochores. Because the CENP-TW complex is one of the most deeply bound kinetochore components at the interface with centromeric chromatin (Fachinetti et al., 2013; Gascoigne et al., 2011; Hori et al., 2013; McKinley et al., 2015; Suzuki et al., 2014; Wan et al., 2009), failure to fully incorporate it in our reconstitutions suggests that an additional binding interface, not necessarily on DNA, may be missing. Characterization of the interaction of CENP-T with the Mis12 complex, on the other hand, brought to light striking similarities with the way that CENP-C interacts with this complex (Petrovic et al., 2016), and offered a glimpse on the potential regulation of this interaction by Aurora B kinase.

**Figure 9.**
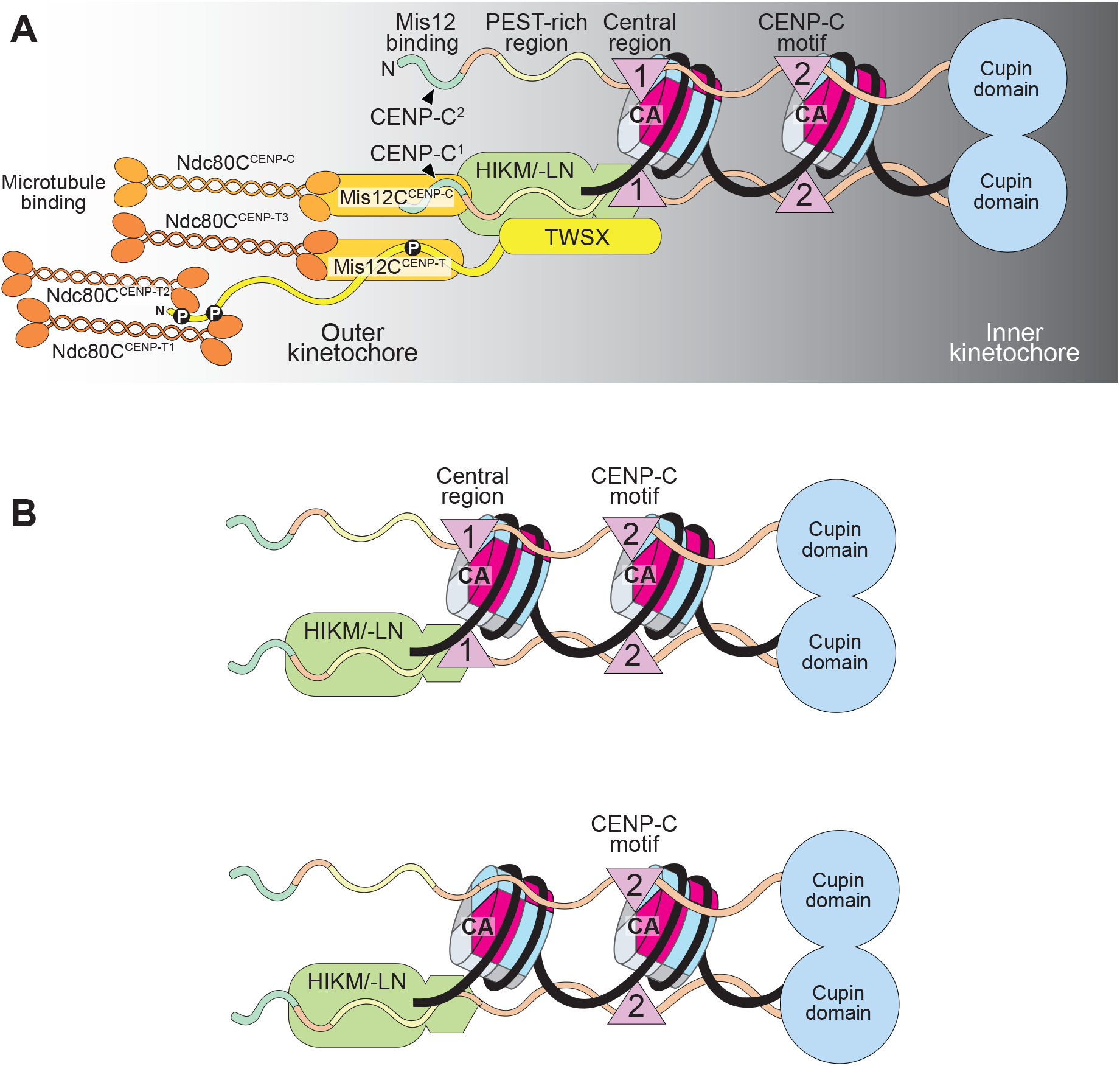
Summary of main conclusions. **A**) The CENP-C dimer binds two nucleosomes (both being represented as CENP-A nucleosomes here). For clarity, binding interactions are only shown for one of the two protomers in the CENP-C dimer. CENP-HIKM/-LN binds near the PEST region, where it attracts CENP-TW. One full KMN network is recruited through the N-terminal region of CENP-C. Another one is recruited through CENP-TW. The latter recruits two additional NDC80 complexes. Therefore, each CENP-C protomer is associated with two KMN network complexes, and a total of four NDC80 complexes. KNL11 was omitted for clarity. **B**) CENP-C with (top) and without (bottom) the CENP-A binding motif in the central region could nevertheless interact with two nucleosomes through the CENP-C motif and the CENP-HIKM/-LN complex.

Recent single-particle cryo-electron microscopy (EM) structural analyses have illuminated the overall organization of the CCAN^Ctf19^ complex of *Saccharamyces cerevisiae* (Hinshaw and Harrison, 2019; Yan et al., 2019), with insights into its “point” single kinetochore module. With the only known exception of kinetoplastids (Akiyoshi and Gull, 2014), eukaryotic kinetochores are evolutionarily related, despite considerable divergence in certain phyla, where loss of one or more CCAN subunits is not uncommon (Drinnenberg and Akiyoshi, 2017; van Hooff et al., 2017). This seems to imply that regional kinetochores are built from multiple, adjacent repetitions of a module related to *Saccharomyces cerevisiae*’s, and that the particle we have reconstituted is, or approaches, the human module. Structural analyses of kinetochores from additional species will have to clarify the extent to which CCAN complexes from other eukaryotes, including humans, resemble the yeast module.

CENP-C is found at kinetochores of most species, likely because its roles in connecting the CENP-A nucleosome to the outer kinetochore and as an organizer of the CCAN complex are indispensable. The apparent co-linearity of binding motifs in CENP-C with the outer-inner kinetochore axis prompted us to define this protein as a blueprint for kinetochore assembly (Klare et al., 2015; Pentakota et al., 2017; Screpanti et al., 2011; Weir et al., 2016). To gain deeper insights into how CENP-C shapes the kinetochore-centromere interface, we harnessed a protein ligation technique (Zakeri et al., 2012) to obtain a full-length version of human CENP-C. Electroporated in cells acutely depleted of endogenous CENP-C, the fusion protein was functional, showcasing the power of electroporation for targeted protein delivery and functional studies with recombinant proteins in cells (Alex et al., 2019).

Our observation that CENP-C^F^ binds selectively to CENP-A nucleosomes over H3 nucleosomes even at micromolecular concentrations *in vitro* was surprising, as previous studies had reported that at least the CENP-C motif, when isolated from the rest of CENP-C, binds relatively non-selectively to CENP-A and H3 NCPs (Ali-Ahmad et al., 2019; Kato et al., 2013), although another study failed to detect an interaction of the CENP-C motif with H3.3 nucleosomes (Arimura et al., 2014). Phosphorylation of CENP-C near the CENP-C motif has been recently shown to facilitate the interaction with CENP-A^NCPs^ *in vitro* and CENP-C kinetochore localization (Watanabe et al., 2019). The interaction of CENP-C with two nucleosomes, implied by the presence of CENP-A^NCP^-binding motifs in the central region and the CENP-C motifs, had been postulated (Kato et al., 2013; Musacchio and Desai, 2017) but remained undemonstrated. Here, we showed that CENP-C interacts with two disconnected CENP-A nucleosomes, or with a reconstituted di-nucleosome. In absence of CCAN proteins, this ability reflects having two CENP-A^NCP^-binding motifs, because mutation of both motifs abrogated binding of CENP-C to CENP-A^NCPs^ *in vitro*, while mutation of either motif reduced the amount of bound CENP-A by half, suggesting that each binding site achieved saturation, at least at the low micromolar concentration of our experiments.

Importantly, many CENP-C orthologues have only the CENP-C motif and no evident CENP-A binding motif in the central motif (Kato et al., 2013; Kral, 2015). Even these CENP-C variants, however, may interact with two nucleosomes, because CENP-C interacts with CENP-A^NCPs^ not only through its CENP-A binding motif(s) but also through CENP-N (Figure 9B). Exemplifying this, addition of CENP-HIKM and CENP-LN complexes to CENP-A^NCP^ binding-assays performed with impairing mutants of the CENP-A^NCPs^ binding motifs of CENP-C rescued binding to CENP-A^NCPs^, while further mutation of the CENP-HIKM and CENP-LN binding sites largely abrogated binding *in vitro* and in cells. CENP-C localized quite robustly to centromeres after mutation or even deletion of both CENP-C motifs, as also reported in other studies (Carroll et al., 2010; Guo et al., 2017; Watanabe et al., 2019). Thus, centromere recruitment of CENP-C relies not only on the CENP-A^NCPs^ binding motifs, but equally, or even preferentially, on interactions with the CCAN. Studies of CENP-C localization in chicken DT40 cells have also demonstrated that recruitment of CENP-C requires CENP-HIKMLN and CENP-A nucleosome binding, but that the dependence on these interactions varies depending on the cell cycle phase (Nagpal et al., 2015; Watanabe et al., 2019). Rapid depletion of CENP-I or CENP-N in human DLD-1 cells demonstrated a partial dependence of CENP-C on these proteins for kinetochore localization in interphase but not in mitosis (McKinley et al., 2015). Other studies in human cells showed partially reduced levels of CENP-C when single CCAN components were depleted by RNAi (Carroll et al., 2009; Cheeseman and Desai, 2008; Weir et al., 2016). Collectively, these analyses identify CENP-C as an organizer of centromeric chromatin and of the inner kinetochore. They also pinpoint that the cell cycle, likely through phosphorylation, influences the localization requirements of CENP-C (Watanabe et al., 2019). Regretfully, structural analyses on yeast Ctf19^CCAN^ have not yet visualized the Mif2p^CENP-C^ polypeptide chain, either because it was not included in the reconstitution (Hinshaw and Harrison, 2019) or because it was for the most part invisible in density maps (Yan et al., 2019), a likely consequence of its predicted intrinsic disorder.

An interesting potential implication of the modular organization of regional kinetochores is that each module, in isolation, may encode its epigenetic specification and propagation. If this were true, it might ultimately become possible to promote CENP-A loading in a reconstituted system provided that the loading machinery is offered the appropriate modular substrate. From a mechanistic perspective, however, how the organization of centromeric chromatin contributes to the recruitment there of specialized CENP-A loading machinery, including the CENP-A chaperone HJURP, the MIS18:M18BP1 complex, and PLK1 kinase, and how CENP-A becomes loaded on chromatin by this machinery remains unclear (Camahort et al., 2007; Dunleavy et al., 2007; Dunleavy et al., 2009; Erhardt et al., 2008; Foltz et al., 2009; Fujita et al., 2007; Hayashi et al., 2004; Maddox et al., 2007; McKinley and Cheeseman, 2014; Mizuguchi et al., 2007; Sanchez-Pulido et al., 2009; Stoler et al., 2007).

Two features of CENP-A nucleosomes that must be considered in this context are 1) that once incorporated into centromeric chromatin, CENP-A is usually retained over extended, sometimes year-long periods of time; and 2) that the levels of CENP-A at each chromosome appear to be maintained intergenerationally, implying that the amounts of newly loaded CENP-A are at least in first approximation identical to those of the centromere-residing CENP-A (Bodor et al., 2013; Falk et al., 2015; Jansen et al., 2007; Raychaudhuri et al., 2012; Smoak et al., 2016). Suppression by cyclin-dependent kinase (CDK) limits loading to the early G1 phase of the cell cycle (Jansen et al., 2007; Pan et al., 2017; Schuh et al., 2007; Silva et al., 2012). The levels of CENP-A then halve during DNA replication, when CENP-A is equally distributed to the sister chromatids. New CENP-A loading is suppressed at this stage, and pre-replication levels of CENP-A are only re-established during the subsequent cell cycle. This contrasts with histone H3.1 deposition, which occurs along with DNA replication (Mendiratta et al., 2019). Equal distribution of CENP-A to the sister chromatids during DNA replication engages HJURP, the same histone chaperone involved in CENP-A deposition, but the mechanistic basis of this process remains unclear (Zasadzinska et al., 2018). Vacancies generated by halving CENP-A may be temporarily filled with the “placeholder” histone H3.3, predicting that when CENP-A is loaded on chromatin in G1, histone H3.3 is concomitantly evicted (Dunleavy et al., 2011).

Collectively, these features suggest a mechanism of CENP-A deposition in which a substrate of existing CENP-A nucleosomes determines how many new CENP-A nucleosomes are generated at every cell cycle. We previously adapted this idea to hypothesize that a minimal unit of two adjacent nucleosomes, with an appropriate complement of CCAN subunits, may satisfy the requirements for a substrate-based CENP-A loading reaction (Musacchio and Desai, 2017). For instance, the placeholder hypothesis (Dunleavy et al., 2011) may be practically implemented on a di-nucleosome scaffold, with a CENP-A nucleosome persisting through the cell cycle and an adjacent nucleosome alternating in composition, with an H3 nucleosome between S-phase and G1, and a CENP-A nucleosome between G1 and S-phase (Musacchio and Desai, 2017). While this remains merely conjectural, dimeric α-satellite units separated by a CENP-B box are a dominant feature of functional human centromeres; CENP-A is greatly enriched in adjacent peaks centered on each α-satellite unit in the dimer, while CENP-T associates more prominently with the linker (Henikoff et al., 2015; Takeuchi et al., 2014; Thakur and Henikoff, 2016; Yoda and Okazaki, 1997).

These considerations emphasize the importance of decoding the localization determinants of the CENP-A loading machinery that limits new CENP-A deposition to these sites, as the dissection of these determinants will reveal the molecular basis of epigenetic centromere specification. These determinants are probably complex and localized through the entire CCAN, because besides CENP-A itself, also CENP-C and CENP-I contribute to centromere recruitment of the Mis18 complex and M18BP1 (Dambacher et al., 2012; French and Straight, 2019; French et al., 2017; Guse et al., 2011; Hori et al., 2017; Kral, 2015; Sandmann et al., 2017; Shono et al., 2015; Stellfox et al., 2016).

All the work of *in vitro* reconstitution with CENP-A or Cse4 (the *S. cerevisiae* ortholog of CENP-A), histone H4, and the H2A/H2B dimer has so far uniformly converged on octameric, left-handed nucleosomes with somewhat looser DNA ends (Black and Cleveland, 2011). Importantly, human kinetochores can be reconstituted on, and bind selectively to, the octameric CENP-A^NCPs^, as shown here and elsewhere (Weir et al., 2016), lending support to the idea that the left-handed octamer is the basic form of centromeric chromatin. CENP-A has been shown to form octameric nucleosomes also on centromeric chromatin in higher eukaryotes (Black and Cleveland, 2011; Dunleavy et al., 2013), but a possible caveat is that extensive nuclease digestions of centromeric chromatin leads to dissociation of CCAN subunits that might trigger post-processing re-organization of CENP-A (Ando et al., 2002; Thakur and Henikoff, 2016).

In *S. cerevisiae*, the single kinetochore “module” is built on a specific, conserved centromere sequence (Fitzgerald-Hayes et al., 1982), and protects ~125 bp of DNA from nuclease digestion, of which approximately 85 bp, sufficient for a single gyre of DNA, wrapped around a CENP-A core proposed to be forming a tetrasome (Cole et al., 2011; Henikoff and Furuyama, 2012; Krassovsky et al., 2012). In addition, the DNA of yeast centromeres is positively supercoiled (Diaz-Ingelmo et al., 2015; Furuyama and Henikoff, 2009; Huang et al., 2011). The recent cryo-EM reconstruction of the Ctf19^CCAN^:Cse4^NCPs^ complex (Cse4 is the *S. cerevisiae* ortholog of CENP-A), however, did not capture these features, as it was obtained with an octameric, left-handed Cse4^NCP^ particle containing the 147-bp Widom 601 DNA sequence (Yan et al., 2019). In this structure, Chl4^CENP-N^ occupies a different position from that previously observed with the isolated CENP-N protein on reconstituted CENP-A^NCPs^ (Chittori et al., 2018; Pentakota et al., 2017; Tian et al., 2018), and that seems hardly compatible with the established function of human CENP-N in decoding the CENP-A L1 loop, where the sequences of CENP-A and H3 diverge (Carroll et al., 2010; Logsdon et al., 2015).

Thus, a fully convincing case for an octameric, left-handed nucleosome in *S. cerevisiae* has not yet been made, and the hypothesis that at least in this organism centromeric chromatin adopts a different organization remains worth of further consideration. In view of the possible modularity of regional kinetochores discussed above and of the considerable conservation of the CCAN subunits, and despite the considerable evidence supporting octameric CENP-A nucleosomes, presence of related non-canonical features in other eukaryotes remains a concrete possibility. How non-canonical features of centromeric chromatin could be built and stabilized, however, remains enigmatic. Clearly, biochemical reconstitution can give the impulse required to solve this conundrum. A benchmark for successful reconstitution could be the reproduction of salient features of the original cellular object, including stoichiometries and successful binding interactions with the CENP-A loading machinery. Our future steps in biochemical reconstitutions will focus on this fundamental and still unresolved question.

## Acknowledgments

We are grateful to Daniele Fachinetti and Don C. Cleveland for sharing reagents, and to Ingrid Hoffmann, Carolin Koerner, Lisa Schulze, Isabelle Stender, Annika Take, Beate Voss, and Sabine Wohlgemuth for general technical assistance, and to Franziska Müller and Petra Janning for help with mass spectrometry experiments. AM gratefully acknowledges funding by the Max Planck Society, the European Research Council (ERC) Advanced Investigator Grant RECEPIANCE (proposal 669686), and the DFG’s Collaborative Research Centre (CRC) 1093.

## Author Contributions

KW: Conceptualization, Investigation, Project administration, Visualization, Writing – Original Draft Preparation; AP: Conceptualization, Investigation, Visualization; DP: Conceptualization, Investigation, Visualization; BH: Investigation, Project administration, Visualization; DV: Investigation; AM: Conceptualization, Funding Acquisition, Project administration, Supervision, Visualization, Writing – Original Draft Preparation.

## Materials and Methods

### Plasmids

Plasmids for the production of *X. laevis* H2A, H2B, H3 and H4 histones were a gift from D. Rhodes. Plasmids for the production of human CENP-A:H4 histone tetramer were a gift from A. F. Straight. Plasmids for the production of the ‘601’ 145-bp DNA were a gift from C. A. Davey. The 181 bp CEN1-like DNA1 sequence and the 205 bp CEN1-like DNA2 sequence were purchased from GeneArt. Eight tandem repeats of CEN1-like DNA fragment 1 or CEN1-like DNA fragment 2 were cloned into a pUC18 plasmid using restriction enzyme-based cloning and Gibson assembly.

The plasmid pETDuet1-mChP-CENPW-HaloT-CENPT-8His was generated by insertion of the coding sequence of mCherry-tagged CENP-W followed by a ribosome binding site and the coding sequence of Halo-tagged CENP-T between *Nco*I and *Xho*I sites of pETDuet-8His (Pan et al., 2017). The plasmids pETDuet1-mChP-CENPW-HaloT-CENPT(1-549)-8His and the plasmid pETDuet1-mChP-CENPW-HaloT-CENPT(1-529)-8His were generated by exchanging the coding sequence of CENP-T with CENP-T 1-549 or CENP-T 1-529 using restriction enzyme-based cloning.

The plasmid pETDuet1-6His-TEV-MBP was generated by subcloning the coding sequence of *E. coli* maltose binding protein with a N-terminal 6His-tag and a TEV protease cleavage site between *Nco*I and *Sal*I sites of pETDuet-1.

A synthetic cDNA segment encoding human CENP-X isoform 1, codon-optimized for expression in bacteria, was subcloned in pET28a using *Xba*I and *Sal*I. A cDNA segment encoding human CENP-S isoform 1 was subcloned into the second cassette of the same plasmid using *Sal*I and *Not*I.

Codon-optimized cDNA of human CENP-C, CENP-T and CENP-W was purchased from GeneArt (Life Technologies). The plasmids containing the SpyCatcher and SpyTag sequences were ordered from Addgene. The plasmid pETDuet-MBP-8His was generated as previously described (Pan et al., 2017). The PCR-amplified CDS of CENP-C-601-943 or CENP-C-721-943 was inserted between the N-terminal MBP-tag and the C-terminal 8His-Tag using *Bam*HI and *Xho*I. The SpyTag sequence was inserted between the N-terminal MBP-tag and the CENP-C-601-943 CDS using *Bgl*2 and *Bam*HI. The PCR-amplified CDS of CENP-C-1-600 was inserted after the N-terminal MBP-tag using *Nhe*I and *Xho*I. The PCR-amplified CDS of SpyCatcher was inserted using *Kpn*I and *Sal*I. In some constructs, MBP was exchanged against mCherry using *Bgl*2 and *Nhe*I. All point mutations and sequence deletions were introduced by PCR-based site-directed mutagenesis.

For the co-expression of EGFP-CENP-C and different mCherry-tagged Cupin constructs, the CDS encoding full-length human CENP-C and three different CDS encoding Cupin domains from human, *S. cerevisiae* and *D. melanogaster* were subcloned in pcDNA5-EGFP-NLS-P2AT2A-mCherry-PTS1 (Pan et al., 2017), a modified version of pcDNA5-FRT/TO (Thermo Fisher Scientific). The CDS of NLS was exchanged against the CDS of human CENP-C-1-943 using *Bam*HI and *Xho*I. The CDS of PTS was exchanged against the CDS of human CENP-C-773-943, *S. cerevisiae* CENP-C-321-529 and *D. melanogaster* CENP-C-1127-1411 using *Nhe*I and *Xma*I. The CDS of *S. cerevisiae* CENP-C-321-529 was PCR-amplified from genomic yeast DNA. The CDS of *D. melanogaster* CENP-C was obtained from GeneArt. For the single expression of EGFP-tagged CENP-C constructs, the CDS of different CENP-C fragments were subcloned in a pcDNA5-EGFP-P2AT2A-2xStrepFLAG plasmid that was generated by inverse PCR. The PCR-amplified CDS of EGFP was inserted using *Kpn*I and *Bam*HI. The PCR-amplified CDS of CENP-C1-600 was inserted using *Hind*III and *Kpn*I. The PCR-amplified CDS of CENP-C-601-943, CENP-C-601-772, CENP-C-721-943, CENP-C-760-943 were inserted using *Bam*HI and *Xho*I. The PCR-amplified CDS of *S. cerevisiae* CENP-C-321-529, *D. melanogaster* CENP-C-1127-1411 and the LacI dimerization domain (LacI-57-339) were inserted using *Nhe*I and *Xma*I.

### Purification of Cen1-like DNA fragments for nucleosome reconstitution

The pUC18 plasmids were transformed into competent *E. coli* cells. Inoculated bacteria cultures were incubated overnight at 37 °C in TB media supplemented with 100 μg/ml Ampicillin. Subsequently the plasmid DNA was purified using a Giga Purification Kit (Macherey-Nagel). After *EcoR*V digestion the insert DNA was separated from the backbone DNA by PEG precipitation. The isolated DNA fragments were digested with *BstX*I/*Bgl*I (DNA1) and *Bgl*I/*Dra*III (DNA2). The digested DNA fragments were loaded on a HiTrap Q FF anion exchange column (GE Healthcare) and eluted with a linear gradient from 0-2000 mM NaCl in 20 bed volumes. Fractions containing the DNA fragments were precipitated with ethanol and dissolved in 2 M NaCl.

### Reconstitution of nucleosomes

CENP-A and H3 containing mononucleosomes assembled on ‘601’ 145-bp DNA were reconstituted precisely as described (Weir et al., 2016). For the assembly of dinucleosomes, CENP-A and H3 containing NCPs were first reconstituted using two different CEN1-like DNA fragments. 1 μM of NCPs reconstituted on CEN1-like DNA fragment 1 and CEN1-like DNA fragment 2 were ligated to each other for 16 h at 4 °C using 8xHis-tagged T4 DNA ligase. The ligated NCPs were incubated with Complete nickel beads (Roche) for 7 h at 4°C to remove the T4 DNA ligase. The ligated dinucleosomes were concentrated, and glycerol was added at a final concentration of 2 % (v/v). The mixture was loaded on a cylindrical gel containing 5 % reduced Bis-acrylamide. Native PAGE was carried out on a Mini Prep Cell apparatus (Bio-Rad) at 1 W constant power. Dinucleosomes were eluted at a constant flow rate of 0.1 mL/min overnight at 4°C into nucleosome buffer (50 mM NaCl, 10 mM triethanolamine (TEA), 1 mM EDTA) and collected in 250 μL fractions on 96-well-plates. The OD260 and OD280 of the individual fractions were measured using a CLARIOstar Plus plater reader (BMG Labtech). Fractions containing dinucleosomes were pooled, concentrated and stored at 4 °C.

### Human cell lines

Parental Flp-In T-REx HeLa cells were a gift from S. Taylor (University of Manchester, Manchester, England, UK). Flp-In T-REx DLD-1 CENP-C-AID-EYFP cells were a generous gift from D. Fachinetti (Institut Curie, Paris, France) and Don C. Cleveland (University of California, San Diego, USA). These cells have both alleles of CENP-C tagged at the C-Terminus with an Auxin-inducible degron (AID)-EYFP fusion (Fachinetti et al., 2015). Furthermore, a gene encoding the plant E3 ubiquitin ligase osTIR1 was stably integrated into the genome of the cells. To induce rapid depletion of the endogenous CENP-C, 500 μM of the synthetic auxin indole-3-acetic acid (IAA) was added to the cells.

### Cell culture

For all cell culture experiments, cells were cultured in Dulbecco’s Modified Eagle Medium (DMEM, PAN Biotech) supplemented with 10 % tetracycline-free fetal bovine serum (FBS, Thermo Fisher), 2 mM Pen/Strep (PAN Biotech) and 2 mM L-Glutamine (PAN Biotech) at 37 °C in a 5 % CO_2_ atmosphere. Expression of CENP-C-GFP fusion proteins was induced by addition of 50 ng/ml doxycycline (Sigma Aldrich) for at least 24 hours.

### CENP-C RNAi

Gene expression of endogenous CENP-C was inhibited using a single siRNA (target sequence: 5’-GGAUCAUCUCAGAAUAGAA-3’ obtained from Sigma-Aldrich), which targets the coding region of CENP-C mRNA. The expression of codon-optimized CENP-C rescue constructs was not affected by the siRNA treatment. 30 nM CENP-C siRNA was transfected using Lipofectamine RNAiMax (ThermoFisher) according to manufacturer’s instructions. To induce the expression of GFP-tagged CENP-C rescue constructs, 50 ng/ml Doxycycline (Sigma) was added to the cells at the time of siRNA transfection. Phenotypes were analyzed 60 hours after transfection by immunofluorescence microscopy or immunoblotting analysis, respectively.

### Generation of stable cell lines

Stable Flp-In T-REx HeLa cell lines constitutively expressing various CENP-C constructs were generated by Flp/FRT recombination. Deletion mutants and point mutants of CENP-C were generated by PCR site-directed mutagenesis, and the sequences of all constructs were verified by Sanger sequencing (Microsynth Seqlab). CENP-C constructs were cloned into a pcDNA5/FRT plasmid and co-transfected with pOG44 (Invitrogen), a plasmid expressing the Flp recombinase, into cells using X-tremeGENE (Roche) according to manufacturer’s instructions. Following 2 weeks of selection in 250 μg ml^−1^ Hygromycin B (Thermo Fisher) and 4 μg ml^−1^ Blasticidin (Thermo Fisher), single cell colonies were collected and subsequently expanded. Expression of the transgenes was checked by immunofluorescence microscopy and Western Blot analysis.

### Immunofluorescence microscopy

The PFA-fixated cells were permeabilized with PBS-T (PBS buffer containing 0.1% Triton X-100) for 10 min and incubated with PBS-T containing 4% BSA for 40 min. Cells were incubated for 90 min at room temperature with CREST/anti-centromere antibody (Antibodies, Inc.; dilution: 1:200 in PBS-T + 2 % BSA), washed three times with PBS-T and were subsequently treated for 30 min with anti-human Alexa 647–conjugated secondary antibody (Jackson ImmunoResearch; dilution: 1:200 in PBS-T + 2 % BSA) to immunostain the centromeres. To visualize DNA, 0.5 μg/ml DAPI (Serva) was added during the last washing step for 3 min. After drying, the coverslips were mounted with Mowiol mounting media (EMD Millipore) on glass slides and imaged using a ×60 oil immersion objective lens on a DeltaVision deconvolution microscope. The Deltavision Elite System (GE Healthcare, UK) is equipped with an IX-71 inverted microscope (Olympus, Japan), a PLAPON 60x/1.42NA objective (Olympus) and a pco.edge sCMOS camera (PCO-TECH Inc., USA). Quantification of centromere signals was performed using the software Fiji with a script for semiautomated processing. Briefly, average projections were made from z-stacks of recorded images. Centromere spots were chosen based on the parameters of shape, size, and intensity using the images of the reference channel obtained with CREST-staining, and their positions were recorded. In the images of the data channels, the mean intensity value of adjacent pixels of a centromere spot was subtracted as background intensity from the mean intensity value of the centromere spot.

### Immunoblotting

HeLa cells were harvested by trypsinization and the cell pellet was washed with PBS. Cells were incubated in lysis buffer (75 mM HEPES pH 7.5, 150 mM NaCl, 1.5 mM EGTA, 10 mM MgCl2, 10% glycerol, 0.1% NP-40, 90 U/ml benzonase (Sigma), 1 mM PMSF and protease inhibitor mix HP Plus (Serva)) for 15 min on ice and finally subjected to sonication using Bioruptor Plus sonication device (Diagenode). The lysed cells were centrifuged at 16,000 x g for 30 min at 4 °C and SDS sample buffer was added to the supernatant. After Tricin-SDS-PAGE, the proteins were blotted on a nitrocellulose membrane and detected by Western Blot analysis. The following primary antibodies were used: Anti-alpha-Tubulin (1:8000, Sigma Aldrich, T9026), anti-CENP-C (1:1000, SI410, gift from S. Trazzi), anti-mCherry (1:2000, Novus Biologicals, NBP1-96752), anti-GFP (HRP-coupled, 1:10,000, Abcam, 190584) and anti-Vinculin (1:10,000, Sigma Aldrich, V9131). As secondary antibodies, we used anti-mouse or anti-rabbit (1:10,000, Amersham, NXA931, NA934) conjugated to horseradish peroxidase. After incubation with ECL western blotting reagent (GE Healthcare), images were acquired with ChemiDocTM MP System (BioRad) using ImageLab 5.1 software.

### Electroporation of recombinant proteins into human cells

To electroporate recombinant mCherry or mCherry-CENP-C protein into DLD-1-CENP-C-AID-EYFP cells and recombinant mCherry-CENP-W/Halo-CENP-T into HeLa cells, the Neon Transfection System Kit (Thermo Fisher) was used. 3 × 10^6^ cells were trypsinized, washed with PBS and resuspended in electroporation Buffer R (Thermo Fisher) to a final volume of 90 μl. Recombinant mCherry-CENP-C and mCherry-CENP-W/Halo-CENP-T protein was diluted 1:2 in buffer R to 15 μM and 30 μl of the mixture was added to the 90 μl cell suspension. After mixing the sample, 100 μl of the mixture was loaded into a Neon Pipette Tip (Thermo Fisher) and electroporated by applying two consecutive 35 ms pulses with an amplitude of 1005 V. The sample was subsequently added to 50 ml of pre-warmed PBS, centrifuged at 500 g for 3 min and trypsinized for 7 min to remove non-internalized extracellular protein. After one additional PBS washing step and centrifugation, the cell pellet was resuspended in DMEM without antibiotics and transferred to a 12-well plate containing poly-L-Lysine coated coverslips. After the electroporation procedure DLD-1-CENP-C-AID-EYFP cells were additionally treated with 500 μM IAA (Sigma) to induce rapid depletion of endogenous CENP-C. Electroporated Halo-CENP-T was labelled using the Halo Tag Oregon Green fluorescent ligand (Promega) according to the manufacturer’s instructions.

### Co-immunoprecipitation experiment

Purified cleared lysates from HeLa Flp-In T-REx cells co-expressing EGFP-tagged CENP-C and mCherry-tagged Cupin fragments from human, *cerevisiae* and *D. melanogaster* were used for co-immunoprecipitation experiments. 4 mg lysate was incubated with 10 μl RFP-Trap magnetic agarose beads (Chromotek) for 3 h at 4 °C. Subsequently, the beads were washed three times with 500 μl washing buffer (75 mM HEPES pH 7.5, 150 mM NaCl, 0.1 % NP-40, 1 x Protease inhibitor cocktail (Serva)). Afterwards, 2 x SDS-PAGE sample loading buffer was added to the dry beads. The samples were boiled for 5 min at 95 °C and analyzed by SDS-PAGE and subsequent Western Blotting analysis.

### Protein expression and purification

The proteins 6xHis-TEV-MBP-CENP-C^1-600^-SpyCatcher, MBP-TEV-SpyTag-CENP-C^601-943^-8xHis, MBP-TEV-CENP-C^721-943^-8xHis, mCherry-PreSc-CENP-W/Halo-TEV-CENP-T-8xHis, 6xHis-PreSc-MBP-CENP-W/Halo-TEV-CENP-T, 6xHis-TEV-CENP-W/Halo-TEV-CENP-T and CENP-X/CENP-S were expressed under same conditions. E. coli BL21(DE3)-Codon-plus-RIL cells transfected with the pETDuet plasmid were grown in 2x YT medium supplemented with Ampicillin and Chloramphenicol at 37°C to an OD600 of 0.8. Then, 0.2 mM ITPG was added to induce protein expression and cells were cultured for 16 hr at 20 °C. Bacterial cells were harvested, washed once with PBS and the cell pellets were stored at −20 °C. The proteins 6xHis-TEV-CENP-W/Halo-TEV-CENP-T and CENP-X/CENP-S were co-expressed by co-transformation of E. coli with two plasmids.

To purify the CENP-C constructs, the bacterial cell pellets were resuspended in lysis buffer (30 mM HEPES pH7.5, 500 mM NaCl, 10 % glycerol, 1mM TCEP) supplemented with 1 mM PMSF and subsequently lysed by sonication. The crude lysate was cleared by centrifugation at 75.000 g for 30 min at 4 °C. The supernatant was passed through a 0.45 μm filter (Sarstedt) and incubated with cOmplete His-Tag purification resin (Roche) for 16 hours on a tube roller (Starlab) at 4 °C. After incubation, the resin was washed with 100 ml lysis buffer A supplemented with 10 mM imidazole. The bound protein was finally eluted in 15 ml lysis buffer supplemented with 400 mM imidazole. The samples were concentrated using centrifugal filters with a 30 kDa mass cut-off (Merck) and were subsequently applied to a Superose 6 10/300 size-exclusion column (GE healthcare) equilibrated in SEC buffer (10 mM HEPES pH 7.5, 300 mM NaCl, 2.5 % glycerol, 1 mM TCEP). SEC was performed under isocratic conditions at a constant flow rate of 0.5 ml/min. 500 μl fractions were collected and relevant fractions were analyzed by SDS-PAGE, pooled, concentrated, flash-frozen and stored at −80 °C until used for further experiments. Purification of mCherry-CENP-W/Halo-CENP-T-8His and 6His-MBP-CENP-W/Halo-CENP-T was identical, but the Lysis buffer contained 1 M NaCl and the SEC buffer contained 500 mM NaCl and 5 % glycerol.

To purify the CENP-TWSX tetramer, the bacterial pellets were resuspended in lysis buffer (50 mM Tris pH 8.0, 1 M NaCl, 10 % glycerol, 1 M EDTA, 5 mM 2-Mercaptoethanol and 1 mM TCEP) supplemented with 1 mM PMSF, lysed by sonication and cleared by centrifugation at 75,000 g at 4°C for 1 hour. The cleared lysate was applied to cOmplete His-Tag purification resin (Roche), pre-equilibrated in lysis buffer and incubated at 4 °C for 6 h on a tube roller. Subsequently, the beads were washed with lysis buffer supplemented with 10 mM imidazole. Subsequently, the CENP-TWSX complex was eluted in lysis buffer supplemented with 400 mM imidazole. His-TEV protease was added and the CENP-TWSX complex was dialysed three times against 2 L of lysis buffer containing a reduced NaCl concentration of 300 mM. After dialysis, the protein complex was loaded on a Hi Trap SP FF column, washed with 10 CV of 15 % Buffer B (50 mM Tris pH 7.4, 2 M NaCl, 5 % glycerol, 1 mM EDTA, 5 mM 2-Mercaptoethanol, 1 mM TCEP) and eluted using a gradient of 300-2000 mM NaCl. The fractions containing CENP-TWSX were pooled, concentrated and loaded onto a Superdex 200 10/300 SEC column (GE Healthcare) pre-equilibrated in SEC buffer (50 mM Tris pH 8.0, 500 mM NaCl, 5 % glycerol, 1 mM TCEP). Fractions containing CENP-TWXS were pooled, concentrated, flash-frozen in liquid nitrogen and stored at −80 °C.

All other proteins were purified as previously described (Huis In ’t Veld et al., 2016; Pentakota et al., 2017; Pesenti et al., 2018; Petrovic et al., 2016; Weir et al., 2016).

### Analytical size-exclusion chromatography

Analytical size-exclusion chromatography (SEC) was performed on a Superose 6 5/150 column (GE Healthcare) in SEC buffer containing 10 mM HEPES pH 7.5, 300 mM NaCl, 2.5 % glycerol and 1 mM TCEP on an ÄKTAmicro system. All samples were eluted under isocratic conditions at 4 °C in SEC buffer at a flow rate of 0.2 ml/min. Elution of proteins was monitored at 280 nm and 254 nm. Elution of mCherry-CENP-C was additionally monitored at 587 nm. 100 μl fractions were collected and analyzed by SDS– PAGE and Coomassie blue staining. To detect complex formation of CENP-C and nucleosomes, the individual proteins were mixed at the indicated concentrations in a total volume of 50 μl, incubated for 30 minutes on ice and finally subjected to SEC.

### SpyTag reaction for in-vitro reconstitution of full-length CENP-C protein

To reconstitute full-length CENP-C, the SpyTag reaction was used (Zakeri et al., 2012). In brief, MBP-TEV-SpyTag-CENP-C^601-943^-8xHis was incubated with TEV protease for 8 h at 4 °C to cleave off the N-Terminal MBP tag. Subsequently, 6xHis-TEV-MBP-CENP-C^1-600^-SpyCatcher was added in 3-fold molar excess. After incubation at 4 °C for 16 h, the reconstituted MBP-CENP-C^1-600^-Spy-CENP-C^601-943^-8xHis protein was separated from unligated protein fragments on a Superose 6 10/300 size-exclusion column (GE Healthcare) as described above. Relevant fractions were pooled and concentrated using 50 kDa cut-off Amicon filters. The concentrated protein was flash frozen and stored at −80 °C.

### Analytical ultracentrifugation

Sedimentation velocity analytical ultracentrifugation was performed in a Beckman XL-A ultracentrifuge at 42,000 rpm at 20 °C. CENP-A NCP (alone and together with MBP-CENP-C^FL^WT or MBP-CENP-CDmotif2) and MBP-CENP-C-721-943 were diluted in AUC buffer (10 mM HEPES pH 7.9, 100 mM NaCl, 2 % glycerol, 1 mM TCEP) and subsequently loaded into standard double-sector centerpieces. The cells were scanned at 260 nm or 280 nm. At least 150 scans were recorded for each sample. Data were analyzed using the program SEDFIT with the model of continuous c(s) distribution (Schuck, 2000). The partial specific volumes of the proteins, buffer density, and buffer viscosity were estimated using the program SEDNTERP. For the DNA of CENP-A NCP, a partial specific volume of 0.55 ml/g was assumed. Figures were generated using the program GUSSI (Brautigam, 2015).

### In vitro phosphorylation by CDK1:Cyclin B

When indicated, samples were phosphorylated during the amylose-resin pull-down experiments using in-house generated CDK1:Cyclin B:CAK1 (CCC). Phosphorylation reactions were set up in binding buffer (10 mM HEPES pH 7.5, 300 mM NaCl, and 1 mM TCEP) containing 100 nM CCC, 3 μM CENP-T/CENP-C/CCAN substrate, 2 mM ATP, and 10 mM MgCl_2_. Reaction mixtures were incubated at 30 °C for 120 min and then used in the pull-down experiments.

### Amylose-resin pull-down assay

The amylose-resin pull-down assay was performed to check the binding of nucleosomes and kinetochore components to MBP-tagged CENP-C variants and MBP-tagged CENP-W/Halo-CENP-T variants. The proteins were diluted with binding buffer (10 mM HEPES pH 7.5, 300 mM NaCl, 2.5% glycerol and 1 mM TCEP) to 3 μM concentration in a total volume of 50 μl and mixed with 20 μl amylose beads (New England BioLabs). After mixing the proteins and the beads, 20 μl were taken as input. The rest of the solution was incubated at 4 °C for 1 h on a thermomixer (Eppendorf) set to 1000 rpm. To separate the proteins bound to the amylose beads from the unbound proteins, the samples were centrifuged at 800 g for 3 min at 4 °C. The supernatant was removed, and the beads were washed four times with 500 μl binding buffer. After the last washing step, 20 μl 2 x SDS-PAGE sample loading buffer was added to the dry beads. The samples were boiled for 5 min at 95 °C and analyzed by SDS-PAGE and CBB staining. In assays performed to investigate the interactions of CENP-T to CCAN the buffer contained 150 mM NaCl, 15 mM HEPES (pH 7.5), 1 mM TCEP and 0.01% Tween20. The samples were incubated for 15 min at 1,400 rpm and 4 °C.

### Binding measurements

Fluorescence polarization experiments were performed with a CLARIOstar plus plate reader using Corning 384 well low volume black round bottom polystyrene NBS microplates. The experiment was performed in 20 μl total reaction volume containing a fixed concentration (20 nM) of the indicated 5-FAM CENP-T or CENP-C peptide (obtained from Genescript). Indicated peptides were mixed in a titration series with an increasing concentration of the respective MIS12C variant in the binding buffer (20 mM Tris pH8, 150 mM NaCl and 1 mM TCEP), and equilibrated for 30 minutes before the measurement at approximately 25 C. Fluorecein (5-FAM) was exited with polarized light at 470 nm, and the emitted light was detected at 525 nm through both horizontal and vertical polarizers. Polarization values are shown as mean +- standard error for the mean for at least 3 replicates. The values of the binding constants were derived through non-linear least square fitting procedure to a single-site binding model using the Origin software.

### Modeling

Multiple sequence alignments were carried out using MAFFT (Katoh et al., 2019). The degree of sequence conservation was calculated using ConSurf (Ashkenazy et al., 2016) and was mapped on the MIS12C structure. Human CENP-T residues 207-221 and 230-250 were modelled on the CENP-C peptide residues 6-22 and 28-48 according to the alignment shown in Figure 6C by fitting the residues into the electron density of the PDB ID code dataset 5LSK structure (MIS12 complex with the CENP-C peptide) using Coot (Emsley and Cowtan, 2004). The side chains with different or insufficient density were modelled according to geometrical constraints. Atomic coordinates of the resulting model are available upon request.

## Supplemental Figure Legends

**Figure S1.**
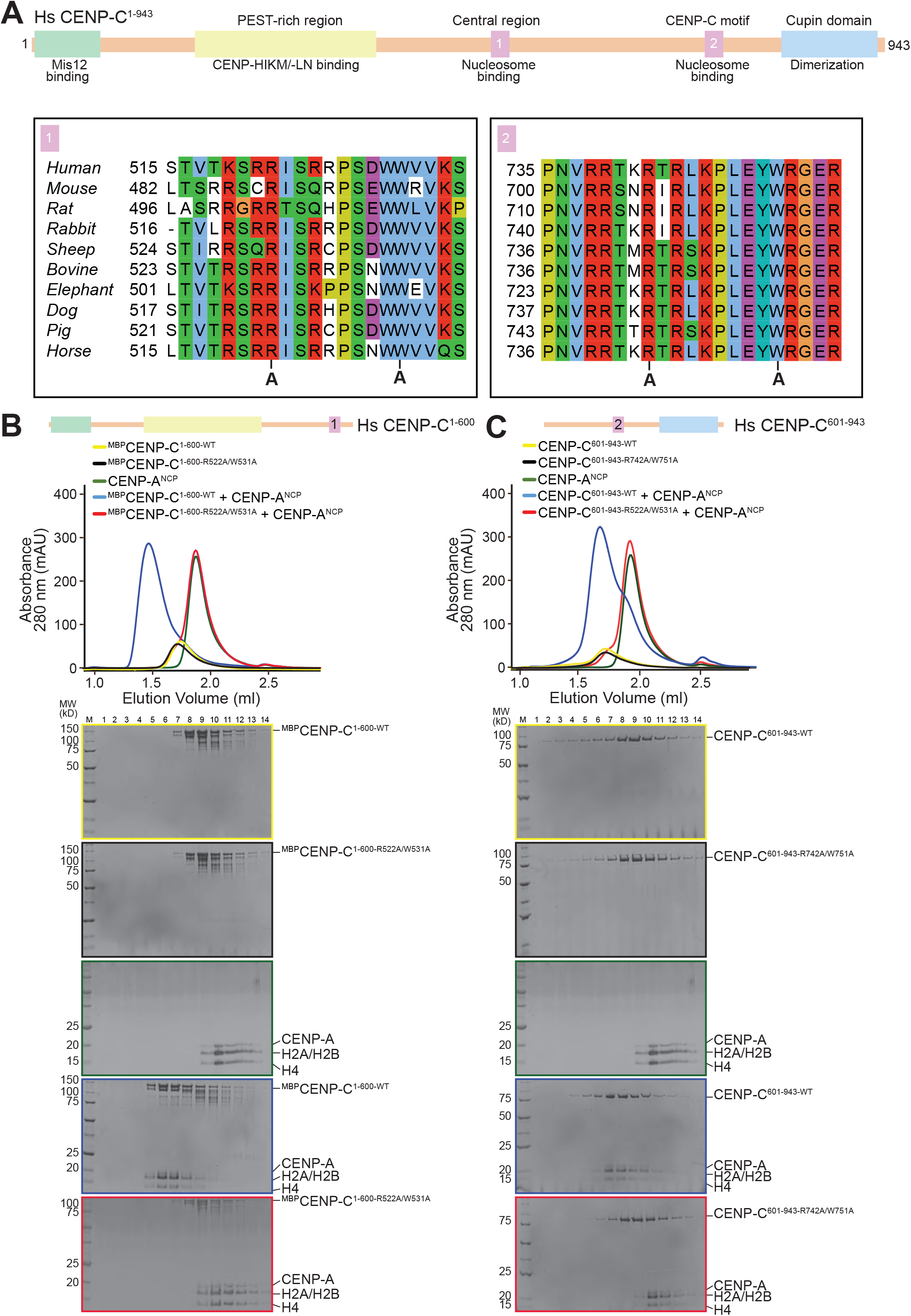
CENP-C central region and CENP-C motif bind CENP-A^NCPs^ individually. **A**) Schematic showing the organization of human CENP-C. Multiple sequence alignments of the central (1) and conserved (2) CENP-C motifs. The ‘A’ below the alignments mark the two residues within each motif which have been substituted by alanine to impair the binding to CENP-A^NCPs^. **B-C**) Analytical SEC experiments using ^MBP^CENP-C^1-600^, ^MBP^CENP-C^601-943^ and CENP-A^NCPs^ to demonstrate that alanine substitutions of two critical residues within each CENP-C motif (Arg522/742 and Trp531/751) prevent the binding of CENP-C to CENP-A^NCPs^.

**Figure S2.**
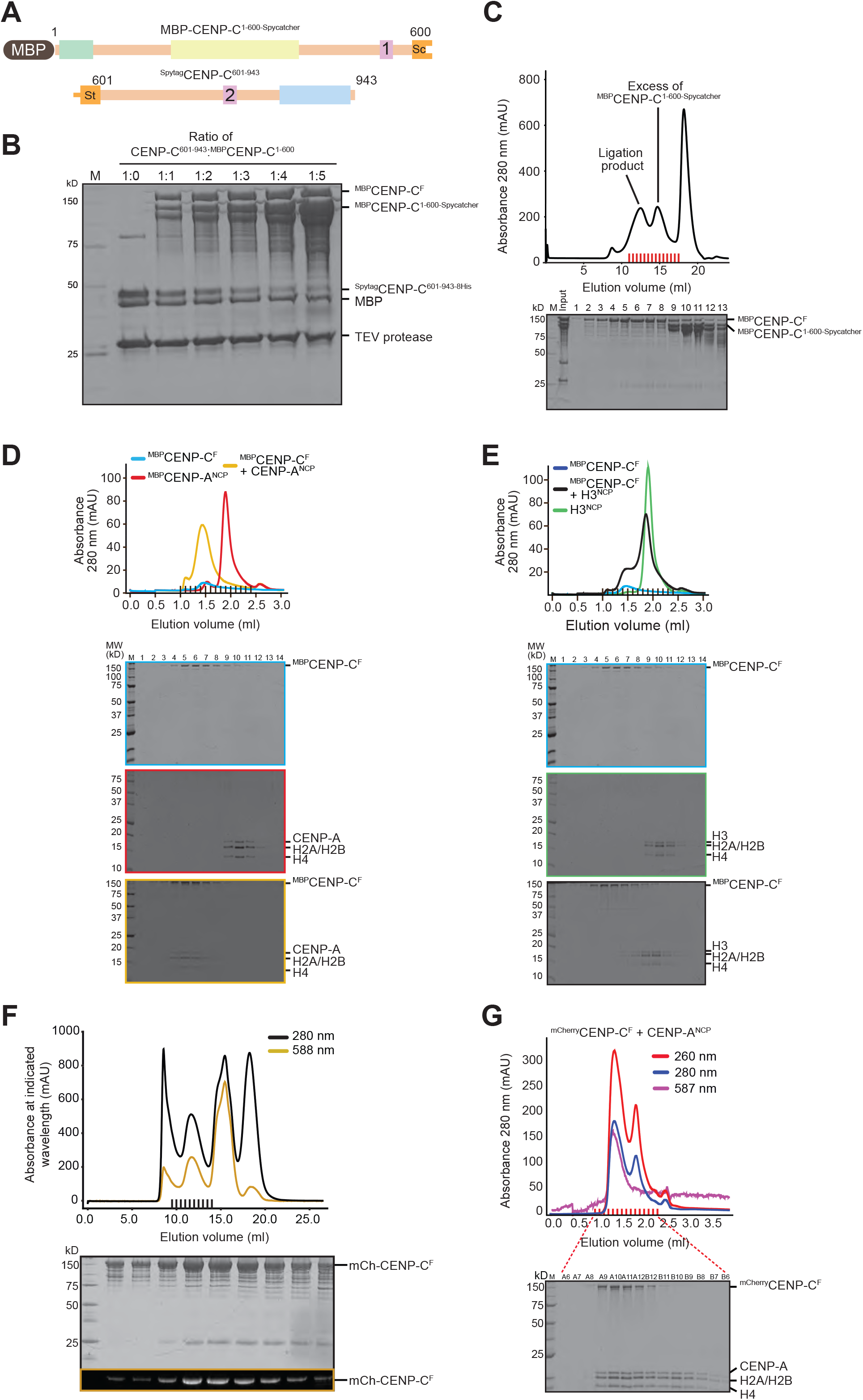
Properties of CENP-C^F^. **A**) Schematic showing the two individual CENP-C fragments that were ligated using the Spycatcher-Spytag system. **B**) SDS-PAGE showing the result of the Spycatcher-Spytag ligation of CENP-C. The ratio of CENP-C^601-943^:CENP-C^1-600^ was varied from 1:0 to 1:5 and is indicated above each lane. **C** Preparative SEC to separate CENP-C^F^ from an excess of unligated CENP-C^1-600-Spycatcher^. The SDS-PAGE result demonstrates that CENP-C^F^ eluted mainly in fractions 2-8, whereas CENP-C^1-600^-SpyCatcher eluted in fractions 9-13. **D-E**) Analytical SEC experiments using ^MBP^CENP-C^F^, CENP-A^NCPs^ and H3^NCPs^ to demonstrate that ^MBP^CENP-C^F^ mainly interacts with CENP-A^NCPs^. **F**) Preparative SEC to purify ^mCherry^CENP-C^F^. Black lines indicate the fractions collected for SDS-PAGE analysis. **G**) Analytical SEC experiments to demonstrate that ^mCherry^CENP-C^F^ interacts with CENP-A NCP. In this experiment, a 2-fold molar excess of CENP-A^NCPs^ was used.

**Figure S3.**
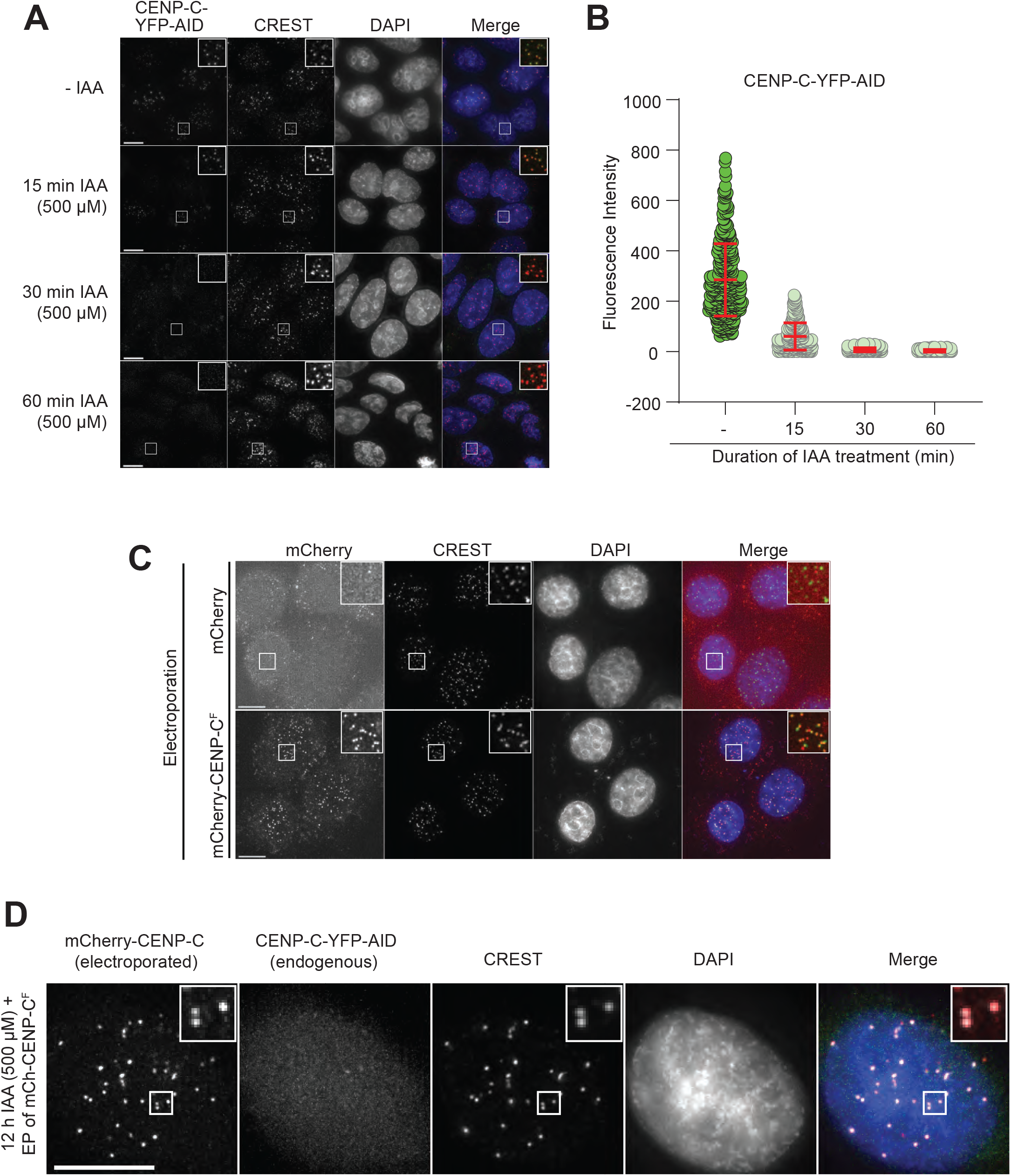
Complementation experiments with CENP-C^F^. **A** Representative images showing the fluorescence of endogenous CENP-C-YFP-AID in fixated DLD-1 cells. Cells were treated with indole acetic acid (IAA) as indicated to rapidly deplete endogenous CENP-C-YFP-AID. Centromeres were visualized by CREST sera and DNA was stained by DAPI. Scale bars indicate 10 μm. **B** Quantification of the CENP-C-YFP intensities. **C** Representative images showing the fluorescence of mCherry and ^mCherry^CENP-C^F^ electroporated into HeLa cells. Centromeres were visualized by CREST sera and DNA was stained by DAPI. Scale bars indicate 10 μm. **D** Representative image showing the fluorescence of ^mCherry^CENP-C^F^ electroporated into DLD-1 cells depleted of endogenous CENP-C-YFP-AID. Centromeres were visualized by CREST sera and DNA was stained by DAPI. Scale bars indicate 10 μm.

**Figure S4.**
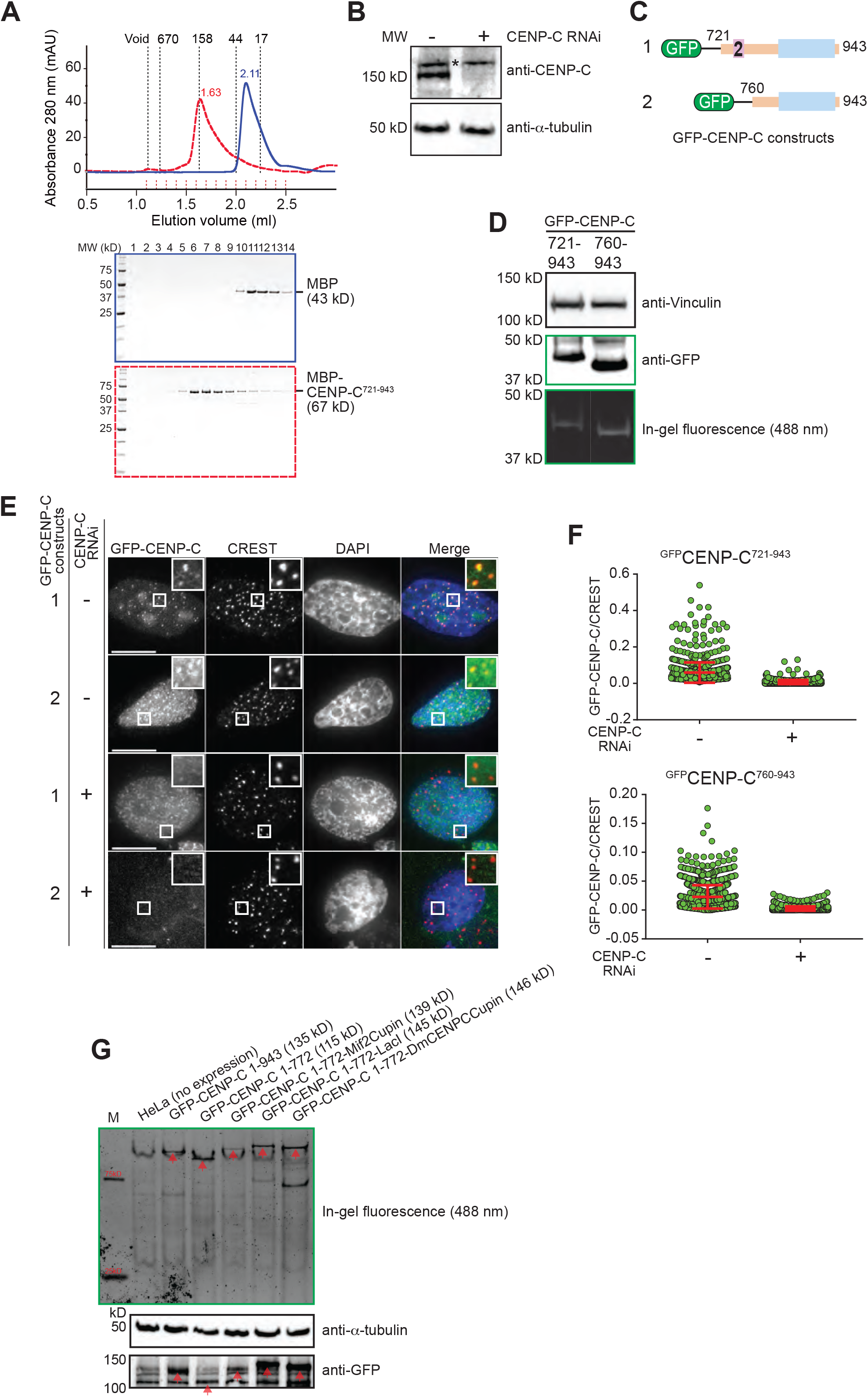
Additional experiments with the CENP-C C-terminal region. **A**) Analytical SEC experiment of MBP and ^MBP^CENP-C^721-943^. Dashed lines represent the elution volume of molecular weight marker proteins ranging from 17 kDa to 670 kDa. **B** Western Blot result showing CENP-C and tubulin. The asterisk marks an unspecific band recognized by the CENP-C antibody. Cells were treated with 30 nM CENP-C siRNA for 60 h. Control cells were treated with RNAiMax without addition of CENP-C siRNA. **C**) The schematic shows the expressed GFP-CENP-C constructs shown in C. **D**) Expression levels of the two indicated CENP-C C-terminal constructs was evaluated by Western blotting and in gel fluorescence. **E**) Representative images showing the fluorescence of two different GFP-CENP-C constructs. The expressed CENP-C construct is indicated at the left of each row. CENP-C RNAi was performed as indicated. Centromeres were visualized by CREST sera and DNA was stained by DAPI. Scale bars indicate 10 μm. **F**) Quantification of the GFP-CENP-C signal intensities. **G**) In-gel fluorescence and anti-GFP western blotting of the indicated constructs used for the experiments in Figure 2G.

**Figure S5.**
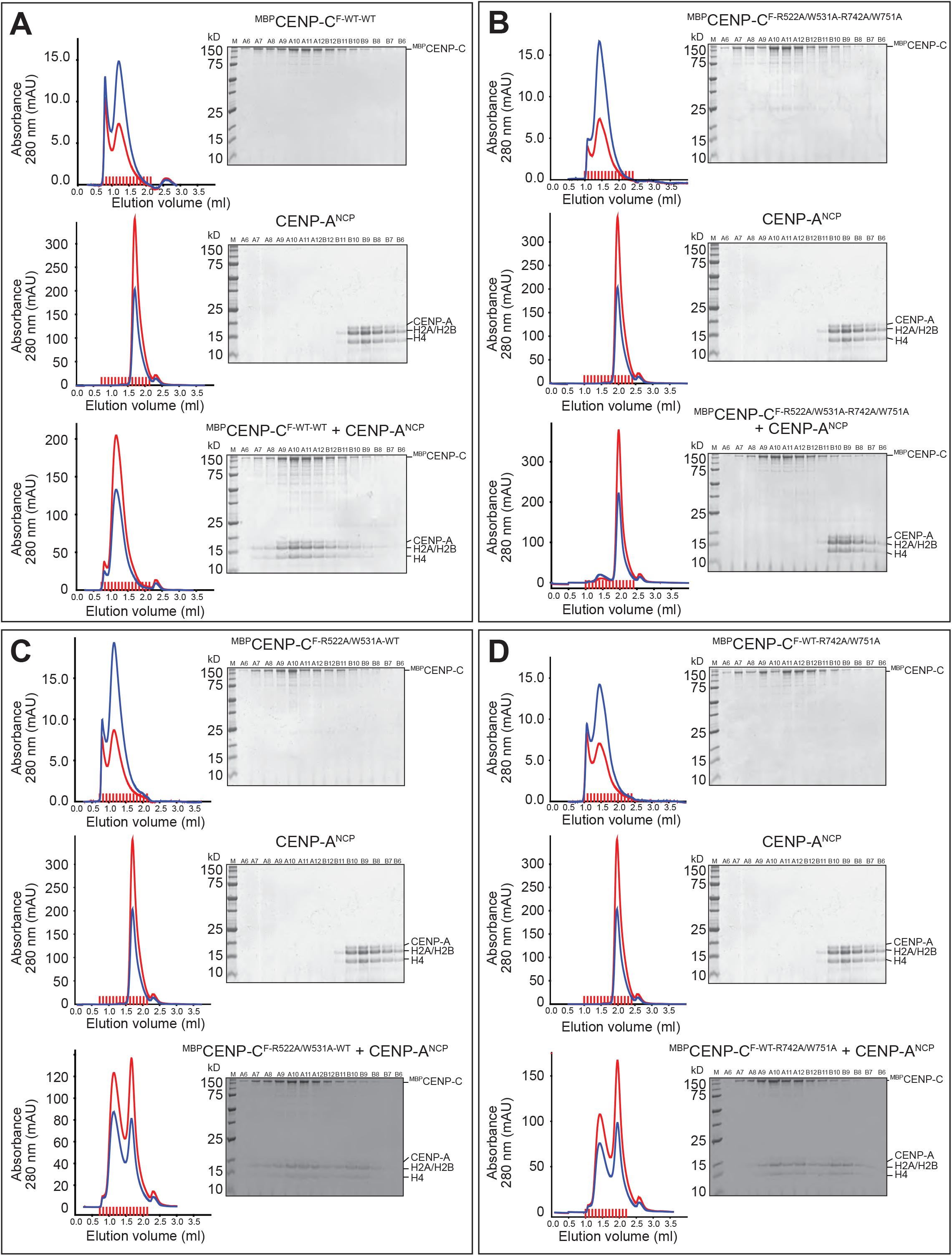
Complete gallery of experiments in Figure 3. **A-D** Analytical SEC experiments of CENP-C^F-WT-WT^, CENP-C^F-R522A/W531A-R742A/W751A^, CENP-C^F-R522A/W531A-WT^, CENP-C^F-WT-R742A/W751A^ and CENP-A^NCPs^ to demonstrate that the interaction between CENP-C and CENP-A NCP depends on functional CENP-C motifs.

**Figure S6.**
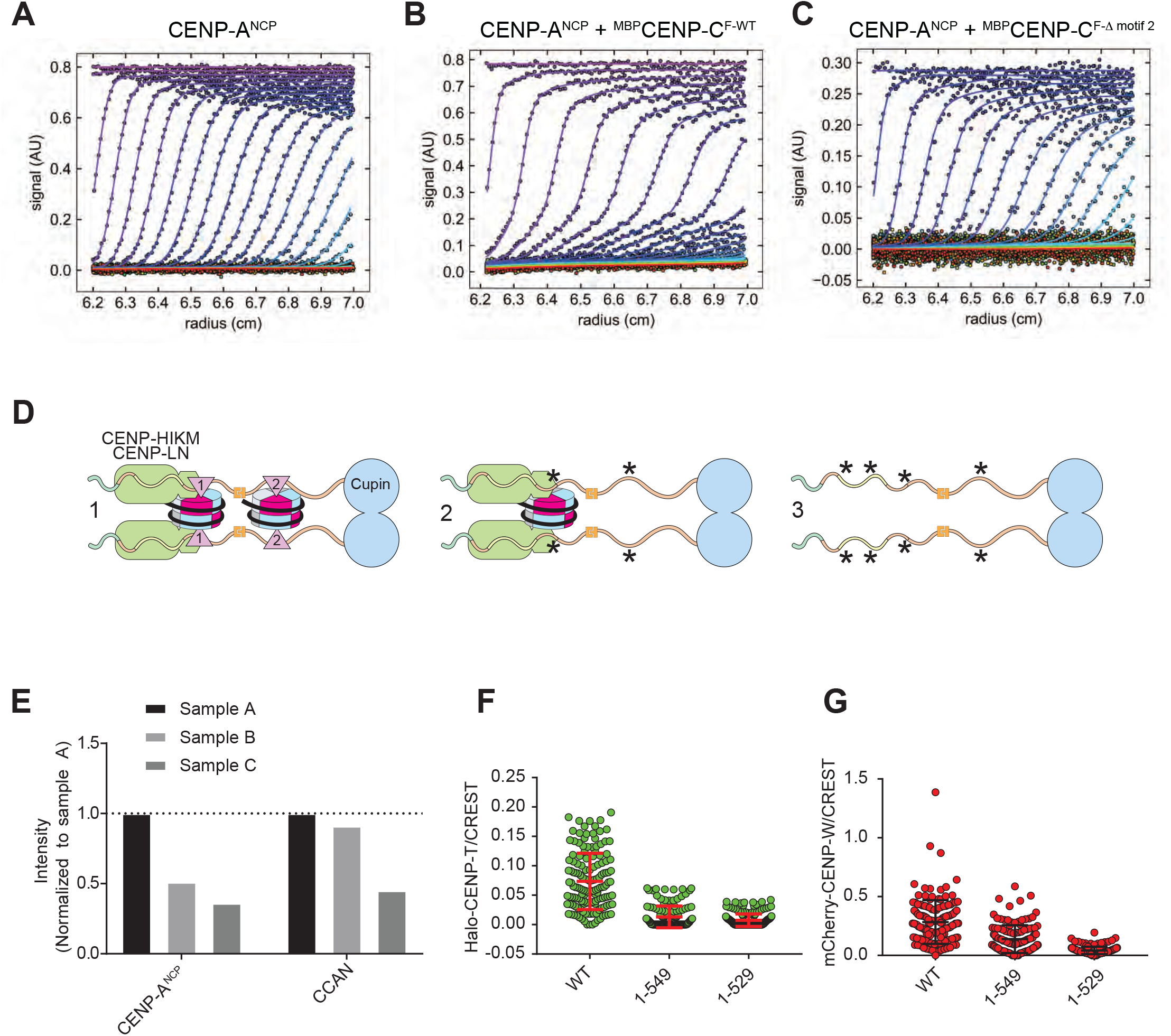
Additional AUC data, band quantifications, and CENP-T localization. **A-C** Best-fitting results of the sedimentation velocity AUC data of CENP-A NCP, CENP-A NCP + MBP-CENP-C^F-WT^ and CENP-A NCP + MBP-CENP-C^F-Dmotif2^. **D** Graphical summary of the pull-down experiment shown in Figure 4A. **E**) Quantification of interaction experiments in Figure 4A. **F-G**) Quantification of localization experiments in Figure 5E.

**Figure S7.**
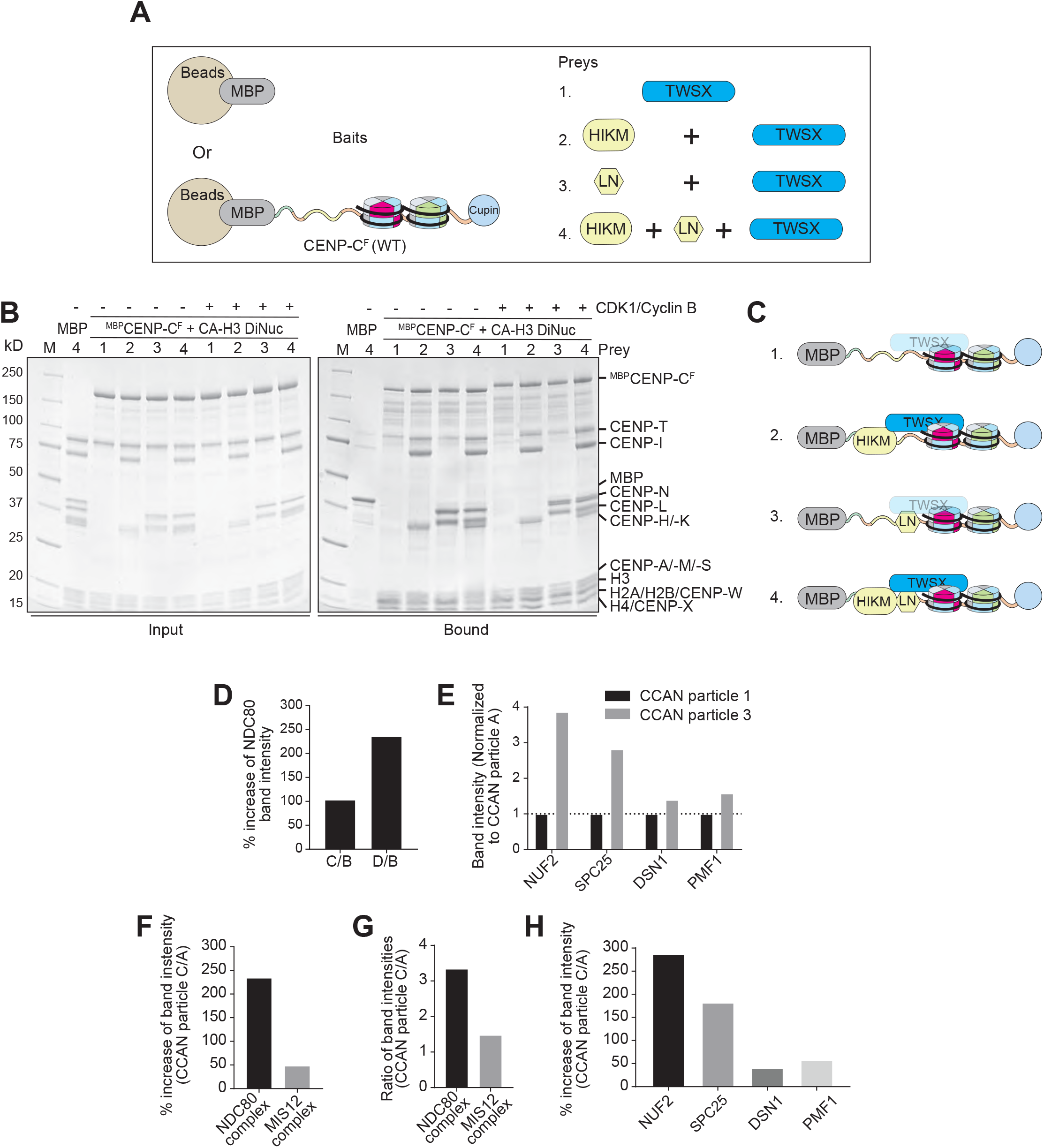
Additional CENP-T:CCAN binding experiments and quantifications. **A** Schematic of the performed amylose-resin pull-down experiment shown in B. MBP or MBP-CENP-C^F^ bound to CENP-A^NCP^/H3^NCP^ dinucleosomes were immobilized on amylose resin as bait. CENP-TWSX, CENP-HIKM and CENP-LN were added as preys as indicated. **B** SDS-PAGE showing the result of the pull-down experiment. The left gel shows the input fractions, the right gel shows the bound fractions. Preys were added as indicated above each lane. **C** Graphical summary of the results shown in B. **D** Quantification showing the increase of the NDC80 band intensities of sample C and D relative to sample B. **E** Quantification showing the individual band intensities of the indicated protein bands shown in Figure 7F. The band intensities were normalized to CCAN particle 1. **F** Quantification showing the % and relative increase of the NDC80 complex and MIS12 complex intensities of CCAN particle 3 relative to CCAN particle 1. The quantified NDC80 complex intensity represents the average of the NUF2 and SPC25 band intensities. The quantified MIS12 complex intensity represents the average of the DSN1 and PMF1 band intensities. **G** Quantification showing the %-increase of the indicated protein bands of CCAN particle 3 relative to CCAN particle 1.

**Table S1.**
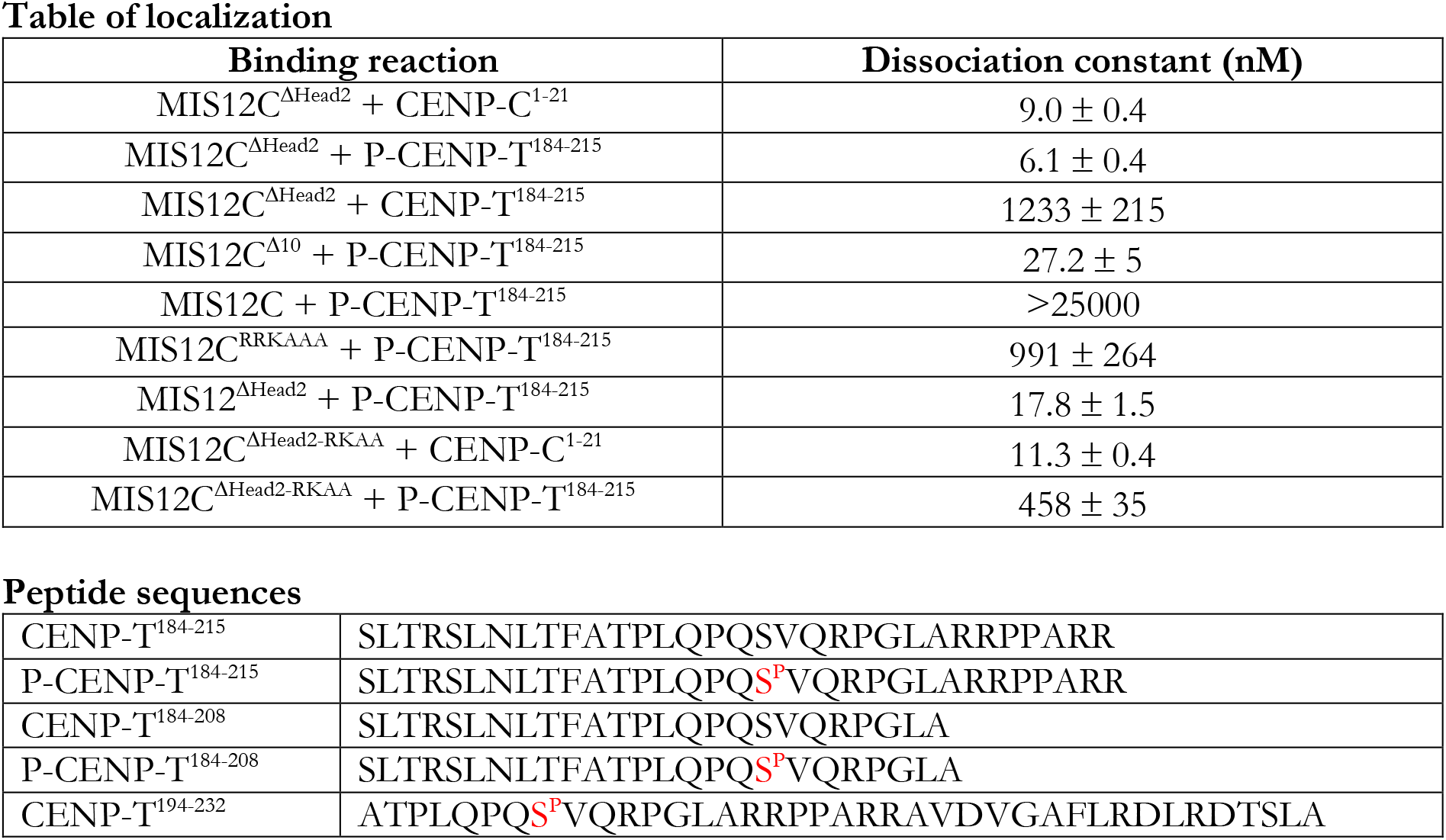
Binding affinities

## References

Akiyoshi, B., and K. Gull. 2014. Discovery of unconventional kinetochores in kinetoplastids. Cell. 156:1247–1258.

Akiyoshi, B., C.R. Nelson, and S. Biggins. 2013. The aurora B kinase promotes inner and outer kinetochore interactions in budding yeast. Genetics. 194:785–789.

Akiyoshi, B., K.K. Sarangapani, A.F. Powers, C.R. Nelson, S.L. Reichow, H. Arellano-Santoyo, Gonen, J.A. Ranish, C.L. Asbury, and S. Biggins. 2010. Tension directly stabilizes reconstituted kinetochore-microtubule attachments. Nature. 468:576–579.

Alex, A., V. Piano, S. Polley, M. Stuiver, S. Voss, G. Ciossani, K. Overlack, B. Voss, S. Wohlgemuth, A. Petrovic, Y. Wu, P. Selenko, A. Musacchio, and S. Maffini. 2019. Electroporated recombinant proteins as tools for in vivo functional complementation, imaging and chemical biology. Elife. 8.

Ali-Ahmad, A., S. Bilokapic, I.B. Schafer, M. Halic, and N. Sekulic. 2019. CENP-C unwraps the human CENP-A nucleosome through the H2A C-terminal tail. EMBO Rep. 20:e48913.

Allis, C.D., and T. Jenuwein. 2016. The molecular hallmarks of epigenetic control. Nat Rev Genet. 17:487–500.

Allu, P.K., J.M. Dawicki-McKenna, T. Van Eeuwen, M. Slavin, M. Braitbard, C. Xu, N. Kalisman, K. Murakami, and B.E. Black. 2019. Structure of the Human Core Centromeric Nucleosome Complex. Curr Biol. 29:2625–2639 e2625.

Ando, S., H. Yang, N. Nozaki, T. Okazaki, and K. Yoda. 2002. CENP-A, -B, and -C chromatin complex that contains the I-type alpha-satellite array constitutes the prekinetochore in HeLa cells. Mol Cell Biol. 22:2229–2241.

Arimura, Y., K. Shirayama, N. Horikoshi, R. Fujita, H. Taguchi, W. Kagawa, T. Fukagawa, G. Almouzni, and H. Kurumizaka. 2014. Crystal structure and stable property of the cancer-associated heterotypic nucleosome containing CENP-A and H3.3. Sci Rep. 4:7115.

Ashkenazy, H., S. Abadi, E. Martz, O. Chay, I. Mayrose, T. Pupko, and N. Ben-Tal. 2016. ConSurf 2016: an improved methodology to estimate and visualize evolutionary conservation in macromolecules. Nucleic Acids Res. 44:W344–350.

Basilico, F., S. Maffini, J.R. Weir, D. Prumbaum, A.M. Rojas, T. Zimniak, A. De Antoni, S. Jeganathan, B. Voss, S. van Gerwen, V. Krenn, L. Massimiliano, A. Valencia, I.R. Vetter, F. Herzog, S. Raunser, S. Pasqualato, and A. Musacchio. 2014. The pseudo GTPase CENP-M drives human kinetochore assembly. Elife. 3:e02978.

Black, B.E., and D.W. Cleveland. 2011. Epigenetic centromere propagation and the nature of CENP-a nucleosomes. Cell. 144:471–479.

Bodor, D.L., J.F. Mata, M. Sergeev, A.F. David, K.J. Salimian, T. Panchenko, D.W. Cleveland, B.E. Black, J.V. Shah, and L.E. Jansen. 2014. The quantitative architecture of centromeric chromatin. Elife. 3:e02137.

Bodor, D.L., L.P. Valente, J.F. Mata, B.E. Black, and L.E. Jansen. 2013. Assembly in G1 phase and long-term stability are unique intrinsic features of CENP-A nucleosomes. Mol Biol Cell. 24:923–932.

Bonner, M.K., J. Haase, J. Swinderman, H. Halas, L.M. Miller Jenkins, and A.E. Kelly. 2019. Enrichment of Aurora B kinase at the inner kinetochore controls outer kinetochore assembly. J Cell Biol. 218:3237–3257.

Brautigam, C.A. 2015. Calculations and Publication-Quality Illustrations for Analytical Ultracentrifugation Data. Methods Enzymol. 562:109–133.

Camahort, R., B. Li, L. Florens, S.K. Swanson, M.P. Washburn, and J.L. Gerton. 2007. Scm3 is essential to recruit the histone h3 variant cse4 to centromeres and to maintain a functional kinetochore. Mol Cell. 26:853–865.

Cao, S., K. Zhou, Z. Zhang, K. Luger, and A.F. Straight. 2018. Constitutive centromere-associated network contacts confer differential stability on CENP-A nucleosomes in vitro and in the cell. Mol Biol Cell. 29:751–762.

Carroll, C.W., K.J. Milks, and A.F. Straight. 2010. Dual recognition of CENP-A nucleosomes is required for centromere assembly. J Cell Biol. 189:1143–1155.

Carroll, C.W., M.C. Silva, K.M. Godek, L.E. Jansen, and A.F. Straight. 2009. Centromere assembly requires the direct recognition of CENP-A nucleosomes by CENP-N. Nat Cell Biol. 11:896–902.

Cheeseman, I.M., J.S. Chappie, E.M. Wilson-Kubalek, and A. Desai. 2006. The conserved KMN network constitutes the core microtubule-binding site of the kinetochore. Cell. 127:983–997.

Cheeseman, I.M., and A. Desai. 2008. Molecular architecture of the kinetochore-microtubule interface. Nat Rev Mol Cell Biol. 9:33–46.

Chik, J.K., V. Moiseeva, P.K. Goel, B.A. Meinen, P. Koldewey, S. An, B.G. Mellone, L. Subramanian, and U.S. Cho. 2019. Structures of CENP-C cupin domains at regional centromeres reveal unique patterns of dimerization and recruitment functions for the inner pocket. J Biol Chem. 294:14119–14134.

Chittori, S., J. Hong, H. Saunders, H. Feng, R. Ghirlando, A.E. Kelly, Y. Bai, and S. Subramaniam. 2018. Structural mechanisms of centromeric nucleosome recognition by the kinetochore protein CENP-N. Science. 359:339–343.

Cohen, R.L., C.W. Espelin, P. De Wulf, P.K. Sorger, S.C. Harrison, and K.T. Simons. 2008. Structural and functional dissection of Mif2p, a conserved DNA-binding kinetochore protein. Mol Biol Cell. 19:4480–4491.

Cole, H.A., B.H. Howard, and D.J. Clark. 2011. The centromeric nucleosome of budding yeast is perfectly positioned and covers the entire centromere. Proc Natl Acad Sci U S A. 108:12687–12692.

Dambacher, S., W. Deng, M. Hahn, D. Sadic, J. Frohlich, A. Nuber, C. Hoischen, S. Diekmann, H. Leonhardt, and G. Schotta. 2012. CENP-C facilitates the recruitment of M18BP1 to centromeric chromatin. Nucleus. 3:101–110.

DeLuca, J.G., W.E. Gall, C. Ciferri, D. Cimini, A. Musacchio, and E.D. Salmon. 2006. Kinetochore microtubule dynamics and attachment stability are regulated by Hec1. Cell. 127:969–982.

Diaz-Ingelmo, O., B. Martinez-Garcia, J. Segura, A. Valdes, and J. Roca. 2015. DNA Topology and Global Architecture of Point Centromeres. Cell Rep. 13:667–677.

Dimitrova, Y.N., S. Jenni, R. Valverde, Y. Khin, and S.C. Harrison. 2016. Structure of the MIND Complex Defines a Regulatory Focus for Yeast Kinetochore Assembly. Cell. 167:1014–1027 e1012.

Drinnenberg, I.A., and B. Akiyoshi. 2017. Evolutionary Lessons from Species with Unique Kinetochores. Prog Mol Subcell Biol. 56:111–138.

Dunleavy, E.M., G. Almouzni, and G.H. Karpen. 2011. H3.3 is deposited at centromeres in S phase as a placeholder for newly assembled CENP-A in G(1) phase. Nucleus. 2:146–157.

Dunleavy, E.M., A.L. Pidoux, M. Monet, C. Bonilla, W. Richardson, G.L. Hamilton, K. Ekwall, P.J. McLaughlin, and R.C. Allshire. 2007. A NASP (N1/N2)-related protein, Sim3, binds CENP-A and is required for its deposition at fission yeast centromeres. Mol Cell. 28:1029–1044.

Dunleavy, E.M., D. Roche, H. Tagami, N. Lacoste, D. Ray-Gallet, Y. Nakamura, Y. Daigo, Y. Nakatani, and G. Almouzni-Pettinotti. 2009. HJURP is a cell-cycle-dependent maintenance and deposition factor of CENP-A at centromeres. Cell. 137:485–497.

Dunleavy, E.M., W. Zhang, and G.H. Karpen. 2013. Solo or doppio: how many CENP-As make a centromeric nucleosome? Nat Struct Mol Biol. 20:648–650.

Dunwell, J.M., A. Culham, C.E. Carter, C.R. Sosa-Aguirre, and P.W. Goodenough. 2001. Evolution of functional diversity in the cupin superfamily. Trends Biochem Sci. 26:740–746.

Earnshaw, W.C., and N. Rothfield. 1985. Identification of a family of human centromere proteins using autoimmune sera from patients with scleroderma. Chromosoma. 91:313–321.

Emsley, P., and K. Cowtan. 2004. Coot: model-building tools for molecular graphics. Acta Crystallogr D Biol Crystallogr. 60:2126–2132.

Erhardt, S., B.G. Mellone, C.M. Betts, W. Zhang, G.H. Karpen, and A.F. Straight. 2008. Genome-wide analysis reveals a cell cycle-dependent mechanism controlling centromere propagation. J Cell Biol. 183:805–818.

Fachinetti, D., H.D. Folco, Y. Nechemia-Arbely, L.P. Valente, K. Nguyen, A.J. Wong, Q. Zhu, A.J. Holland, A. Desai, L.E. Jansen, and D.W. Cleveland. 2013. A two-step mechanism for epigenetic specification of centromere identity and function. Nat Cell Biol. 15:1056–1066.

Fachinetti, D., J.S. Han, M.A. McMahon, P. Ly, A. Abdullah, A.J. Wong, and D.W. Cleveland. 2015. DNA Sequence-Specific Binding of CENP-B Enhances the Fidelity of Human Centromere Function. Dev Cell. 33:314–327.

Falk, S.J., L.Y. Guo, N. Sekulic, E.M. Smoak, T. Mani, G.A. Logsdon, K. Gupta, L.E. Jansen, G.D. Van Duyne, S.A. Vinogradov, M.A. Lampson, and B.E. Black. 2015. Chromosomes. CENP-C reshapes and stabilizes CENP-A nucleosomes at the centromere. Science. 348:699–703.

Fitzgerald-Hayes, M., L. Clarke, and J. Carbon. 1982. Nucleotide sequence comparisons and functional analysis of yeast centromere DNAs. Cell. 29:235–244.

Foltz, D.R., L.E. Jansen, A.O. Bailey, J.R. Yates, 3rd, E.A. Bassett, S. Wood, B.E. Black, and D.W. Cleveland. 2009. Centromere-specific assembly of CENP-a nucleosomes is mediated by HJURP. Cell. 137:472–484.

Foltz, D.R., L.E. Jansen, B.E. Black, A.O. Bailey, J.R. Yates, 3rd, and D.W. Cleveland. 2006. The human CENP-A centromeric nucleosome-associated complex. Nat Cell Biol. 8:458–469.

French, B.T., and A.F. Straight. 2019. CDK phosphorylation of Xenopus laevis M18BP1 promotes its metaphase centromere localization. EMBO J.

French, B.T., F.G. Westhorpe, C. Limouse, and A.F. Straight. 2017. Xenopus laevis M18BP1 Directly Binds Existing CENP-A Nucleosomes to Promote Centromeric Chromatin Assembly. Dev Cell. 42:190–199 e110.

Fujita, Y., T. Hayashi, T. Kiyomitsu, Y. Toyoda, A. Kokubu, C. Obuse, and M. Yanagida. 2007. Priming of centromere for CENP-A recruitment by human hMis18alpha, hMis18beta, and M18BP1. Dev Cell. 12:17–30.

Fukagawa, T., and W.C. Earnshaw. 2014. The centromere: chromatin foundation for the kinetochore machinery. Dev Cell. 30:496–508.

Furuyama, T., and S. Henikoff. 2009. Centromeric nucleosomes induce positive DNA supercoils. Cell. 138:104–113.

Gascoigne, K.E., K. Takeuchi, A. Suzuki, T. Hori, T. Fukagawa, and I.M. Cheeseman. 2011. Induced ectopic kinetochore assembly bypasses the requirement for CENP-A nucleosomes. Cell. 145:410–422.

Guo, L.Y., P.K. Allu, L. Zandarashvili, K.L. McKinley, N. Sekulic, J.M. Dawicki-McKenna, D. Fachinetti, G.A. Logsdon, R.M. Jamiolkowski, D.W. Cleveland, I.M. Cheeseman, and B.E. Black. 2017. Centromeres are maintained by fastening CENP-A to DNA and directing an arginine anchor-dependent nucleosome transition. Nat Commun. 8:15775.

Guse, A., C.W. Carroll, B. Moree, C.J. Fuller, and A.F. Straight. 2011. In vitro centromere and kinetochore assembly on defined chromatin templates. Nature. 477:354–358.

Hara, M., M. Ariyoshi, E.I. Okumura, T. Hori, and T. Fukagawa. 2018. Multiple phosphorylations control recruitment of the KMN network onto kinetochores. Nat Cell Biol. 20:1378–1388.

Hayashi, T., Y. Fujita, O. Iwasaki, Y. Adachi, K. Takahashi, and M. Yanagida. 2004. Mis16 and Mis18 are required for CENP-A loading and histone deacetylation at centromeres. Cell. 118:715–729.

Heeger, S., O. Leismann, R. Schittenhelm, O. Schraidt, S. Heidmann, and C.F. Lehner. 2005. Genetic interactions of separase regulatory subunits reveal the diverged Drosophila Cenp-C homolog. Genes Dev. 19:2041–2053.

Henikoff, J.G., J. Thakur, S. Kasinathan and S. Henikoff. 2015. A unique chromatin complex occupies young alpha-satellite arrays of human centromeres. Sci Adv. 1.

Henikoff, S., and T. Furuyama. 2012. The unconventional structure of centromeric nucleosomes. Chromosoma. 121:341–352.

Hinshaw, S.M., and S.C. Harrison. 2013. An Iml3-Chl4 heterodimer links the core centromere to factors required for accurate chromosome segregation. Cell Rep. 5:29–36.

Hinshaw, S.M., and S.C. Harrison. 2019. The structure of the Ctf19c/CCAN from budding yeast. Elife. 8.

Hinshaw, S.M., and S.C. Harrison. 2020. The Structural Basis for Kinetochore Stabilization by Cnn1/CENP-T. Curr Biol. 30:3425–3431 e3423.

Holland, A.J., D. Fachinetti, J.S. Han, and D.W. Cleveland. 2012. Inducible, reversible system for the rapid and complete degradation of proteins in mammalian cells. Proc Natl Acad Sci U S A. 109:E3350–3357.

Hori, T., M. Amano, A. Suzuki, C.B. Backer, J.P. Welburn, Y. Dong, B.F. McEwen, W.H. Shang, E. Suzuki, K. Okawa, I.M. Cheeseman, and T. Fukagawa. 2008. CCAN makes multiple contacts with centromeric DNA to provide distinct pathways to the outer kinetochore. Cell. 135:1039–1052.

Hori, T., W.H. Shang, M. Hara, M. Ariyoshi, Y. Arimura, R. Fujita, H. Kurumizaka, and T. Fukagawa. 2017. Association of M18BP1/KNL2 with CENP-A Nucleosome Is Essential for Centromere Formation in Non-mammalian Vertebrates. Dev Cell. 42:181–189 e183.

Hori, T., W.H. Shang, K. Takeuchi, and T. Fukagawa. 2013. The CCAN recruits CENP-A to the centromere and forms the structural core for kinetochore assembly. J Cell Biol. 200:45–60.

Huang, C.C., K.M. Chang, H. Cui, and M. Jayaram. 2011. Histone H3-variant Cse4-induced positive DNA supercoiling in the yeast plasmid has implications for a plasmid origin of a chromosome centromere. Proc Natl Acad Sci U S A. 108:13671–13676.

Huis In ’t Veld, P.J., S. Jeganathan, A. Petrovic, P. Singh, J. John, V. Krenn, F. Weissmann, T. Bange, and A. Musacchio. 2016. Molecular basis of outer kinetochore assembly on CENP-T. Elife. 5.

Huis In ’t Veld, P.J., V.A. Volkov, I.D. Stender, A. Musacchio, and M. Dogterom. 2019. Molecular determinants of the Ska-Ndc80 interaction and their influence on microtubule tracking and force-coupling. Elife. 8.

Izuta, H., M. Ikeno, N. Suzuki, T. Tomonaga, N. Nozaki, C. Obuse, Y. Kisu, N. Goshima, F. Nomura, N. Nomura, and K. Yoda. 2006. Comprehensive analysis of the ICEN (Interphase Centromere Complex) components enriched in the CENP-A chromatin of human cells. Genes Cells. 11:673–684.

Jansen, L.E., B.E. Black, D.R. Foltz, and D.W. Cleveland. 2007. Propagation of centromeric chromatin requires exit from mitosis. J Cell Biol. 176:795–805.

Joglekar, A.P., K. Bloom, and E.D. Salmon. 2009. In vivo protein architecture of the eukaryotic kinetochore with nanometer scale accuracy. Curr Biol. 19:694–699.

Joglekar, A.P., D. Bouck, K. Finley, X. Liu, Y. Wan, J. Berman, X. He, E.D. Salmon, and K.S. Bloom. 2008. Molecular architecture of the kinetochore-microtubule attachment site is conserved between point and regional centromeres. J Cell Biol. 181:587–594.

Joglekar, A.P., D.C. Bouck, J.N. Molk, K.S. Bloom, and E.D. Salmon. 2006. Molecular architecture of a kinetochore-microtubule attachment site. Nat Cell Biol. 8:581–585.

Karpen, G.H., and R.C. Allshire. 1997. The case for epigenetic effects on centromere identity and function. Trends Genet. 13:489–496.

Kato, H., J. Jiang, B.R. Zhou, M. Rozendaal, H. Feng, R. Ghirlando, T.S. Xiao, A.F. Straight, and Y. Bai. 2013. A conserved mechanism for centromeric nucleosome recognition by centromere protein CENP-C. Science. 340:1110–1113.

Katoh, K., J. Rozewicki, and K.D. Yamada. 2019. MAFFT online service: multiple sequence alignment, interactive sequence choice and visualization. Brief Bioinform. 20:1160–1166.

Killinger, K., M. Bohm, P. Steinbach, G. Hagemann, M. Bluggel, K. Janen, S. Hohoff, P. Bayer, F. Herzog, and S. Westermann. 2020. Auto-inhibition of Mif2/CENP-C ensures centromere-dependent kinetochore assembly in budding yeast. EMBO J. 39:e102938.

Kim, S., and H. Yu. 2015. Multiple assembly mechanisms anchor the KMN spindle checkpoint platform at human mitotic kinetochores. J Cell Biol. 208:181–196.

Klare, K., J.R. Weir, F. Basilico, T. Zimniak, L. Massimiliano, N. Ludwigs, F. Herzog, and A. Musacchio. 2015. CENP-C is a blueprint for constitutive centromere-associated network assembly within human kinetochores. J Cell Biol. 210:11–22.

Kral, L. 2015. Possible identification of CENP-C in fish and the presence of the CENP-C motif in M18BP1 of vertebrates. F1000Res. 4:474.

Krassovsky, K., J.G. Henikoff, and S. Henikoff. 2012. Tripartite organization of centromeric chromatin in budding yeast. Proc Natl Acad Sci U S A. 109:243–248.

Lang, J., A. Barber, and S. Biggins. 2018. An assay for de novo kinetochore assembly reveals a key role for the CENP-T pathway in budding yeast. Elife. 7.

Logsdon, G.A., E.J. Barrey, E.A. Bassett, J.E. DeNizio, L.Y. Guo, T. Panchenko, J.M. Dawicki-McKenna, P. Heun, and B.E. Black. 2015. Both tails and the centromere targeting domain of CENP-A are required for centromere establishment. J Cell Biol. 208:521–531.

Maddox, P.S., F. Hyndman, J. Monen, K. Oegema, and A. Desai. 2007. Functional genomics identifies a Myb domain-containing protein family required for assembly of CENP-A chromatin. J Cell Biol. 176:757–763.

Malvezzi, F., G. Litos, A. Schleiffer, A. Heuck, K. Mechtler, T. Clausen, and S. Westermann. 2013. A structural basis for kinetochore recruitment of the Ndc80 complex via two distinct centromere receptors. EMBO J. 32:409–423.

McKinley, K.L., and I.M. Cheeseman. 2014. Polo-like kinase 1 licenses CENP-A deposition at centromeres. Cell. 158:397–411.

McKinley, K.L., and I.M. Cheeseman. 2016. The molecular basis for centromere identity and function. Nat Rev Mol Cell Biol. 17:16–29.

McKinley, K.L., N. Sekulic, L.Y. Guo, T. Tsinman, B.E. Black, and I.M. Cheeseman. 2015. The CENP-L-N Complex Forms a Critical Node in an Integrated Meshwork of Interactions at the Centromere-Kinetochore Interface. Mol Cell. 60:886–898.

Medina-Pritchard, B., V. Lazou, J. Zou, O. Byron, M.A. Abad, J. Rappsilber, P. Heun, and A.A. Jeyaprakash. 2020. Structural basis for centromere maintenance by Drosophila CENP-A chaperone CAL1. EMBO J. 39:e103234.

Mendiratta, S., A. Gatto, and G. Almouzni. 2019. Histone supply: Multitiered regulation ensures chromatin dynamics throughout the cell cycle. J Cell Biol. 218:39–54.

Milks, K.J., B. Moree, and A.F. Straight. 2009. Dissection of CENP-C-directed centromere and kinetochore assembly. Mol Biol Cell. 20:4246–4255.

Mizuguchi, G., H. Xiao, J. Wisniewski, M.M. Smith, and C. Wu. 2007. Nonhistone Scm3 and histones CenH3-H4 assemble the core of centromere-specific nucleosomes. Cell. 129:1153–1164.

Musacchio, A., and A. Desai. 2017. A Molecular View of Kinetochore Assembly and Function. Biology (Basel). 6.

Nagpal, H., T. Hori, A. Furukawa, K. Sugase, A. Osakabe, H. Kurumizaka, and T. Fukagawa. 2015. Dynamic changes in CCAN organization through CENP-C during cell-cycle progression. Mol Biol Cell. 26:3768–3776.

Nishimura, K., T. Fukagawa, H. Takisawa, T. Kakimoto, and M. Kanemaki. 2009. An auxin-based degron system for the rapid depletion of proteins in nonplant cells. Nat Methods. 6:917–922.

Nishino, T., F. Rago, T. Hori, K. Tomii, I.M. Cheeseman, and T. Fukagawa. 2013. CENP-T provides a structural platform for outer kinetochore assembly. EMBO J. 32:424–436.

Obuse, C., H. Yang, N. Nozaki, S. Goto, T. Okazaki, and K. Yoda. 2004. Proteomics analysis of the centromere complex from HeLa interphase cells: UV-damaged DNA binding protein 1 (DDB-1) is a component of the CEN-complex, while BMI-1 is transiently co-localized with the centromeric region in interphase. Genes Cells. 9:105–120.

Okada, M., I.M. Cheeseman, T. Hori, K. Okawa, I.X. McLeod, J.R. Yates, 3rd, A. Desai, and T. Fukagawa. 2006. The CENP-H-I complex is required for the efficient incorporation of newly synthesized CENP-A into centromeres. Nat Cell Biol. 8:446–457.

Pan, D., K. Klare, A. Petrovic, A. Take, K. Walstein, P. Singh, A. Rondelet, A.W. Bird, and A. Musacchio. 2017. CDK-regulated dimerization of M18BP1 on a Mis18 hexamer is necessary for CENP-A loading. Elife. 6.

Pekgoz Altunkaya, G., F. Malvezzi, Z. Demianova, T. Zimniak, G. Litos, F. Weissmann, K. Mechtler, F. Herzog, and S. Westermann. 2016. CCAN Assembly Configures Composite Binding Interfaces to Promote Cross-Linking of Ndc80 Complexes at the Kinetochore. Curr Biol. 26:2370–2378.

Pentakota, S., K. Zhou, C. Smith, S. Maffini, A. Petrovic, G.P. Morgan, J.R. Weir, I.R. Vetter, A. Musacchio, and K. Luger. 2017. Decoding the centromeric nucleosome through CENP-N. Elife. 6.

Pesenti, M.E., D. Prumbaum, P. Auckland, C.M. Smith, A.C. Faesen, A. Petrovic, M. Erent, S. Maffini, S. Pentakota, J.R. Weir, Y.C. Lin, S. Raunser, A.D. McAinsh, and A. Musacchio. 2018. Reconstitution of a 26-Subunit Human Kinetochore Reveals Cooperative Microtubule Binding by CENP-OPQUR and NDC80. Mol Cell. 71:923–939 e910.

Petrovic, A., J. Keller, Y. Liu, K. Overlack, J. John, Y.N. Dimitrova, S. Jenni, S. van Gerwen, P. Stege, S. Wohlgemuth, P. Rombaut, F. Herzog, S.C. Harrison, I.R. Vetter, and A. Musacchio. 2016. Structure of the MIS12 Complex and Molecular Basis of Its Interaction with CENP-C at Human Kinetochores. Cell. 167:1028–1040 e1015.

Przewloka, M.R., Z. Venkei, V.M. Bolanos-Garcia, J. Debski, M. Dadlez, and D.M. Glover. 2011. CENP-C is a structural platform for kinetochore assembly. Curr Biol. 21:399–405.

Rago, F., K.E. Gascoigne, and I.M. Cheeseman. 2015. Distinct organization and regulation of the outer kinetochore KMN network downstream of CENP-C and CENP-T. Curr Biol. 25:671–677.

Raychaudhuri, N., R. Dubruille, G.A. Orsi, H.C. Bagheri, B. Loppin, and C.F. Lehner. 2012. Transgenerational propagation and quantitative maintenance of paternal centromeres depends on Cid/Cenp-A presence in Drosophila sperm. PLoS Biol. 10:e1001434.

Sanchez-Pulido, L., A.L. Pidoux, C.P. Ponting, and R.C. Allshire. 2009. Common ancestry of the CENP-A chaperones Scm3 and HJURP. Cell. 137:1173–1174.

Sandmann, M., P. Talbert, D. Demidov, M. Kuhlmann, T. Rutten, U. Conrad, and I. Lermontova. 2017. Targeting of Arabidopsis KNL2 to Centromeres Depends on the Conserved CENPC-k Motif in Its C Terminus. Plant Cell. 29:144–155.

Schleiffer, A., M. Maier, G. Litos, F. Lampert, P. Hornung, K. Mechtler, and S. Westermann. 2012. CENP-T proteins are conserved centromere receptors of the Ndc80 complex. Nat Cell Biol. 14:604–613.

Schuck, P. 2000. Size-distribution analysis of macromolecules by sedimentation velocity ultracentrifugation and lamm equation modeling. Biophys J. 78:1606–1619.

Schuh, M., C.F. Lehner, and S. Heidmann. 2007. Incorporation of Drosophila CID/CENP-A and CENP-C into centromeres during early embryonic anaphase. Curr Biol. 17:237–243.

Screpanti, E., A. De Antoni, G.M. Alushin, A. Petrovic, T. Melis, E. Nogales, and A. Musacchio. 2011. Direct binding of Cenp-C to the Mis12 complex joins the inner and outer kinetochore. Curr Biol. 21:391–398.

Shono, N., J. Ohzeki, K. Otake, N.M. Martins, T. Nagase, H. Kimura, V. Larionov, W.C. Earnshaw, and H. Masumoto. 2015. CENP-C and CENP-I are key connecting factors for kinetochore and CENP-A assembly. J Cell Sci. 128:4572–4587.

Silva, M.C., D.L. Bodor, M.E. Stellfox, N.M. Martins, H. Hochegger, D.R. Foltz, and L.E. Jansen. 2012. Cdk activity couples epigenetic centromere inheritance to cell cycle progression. Dev Cell. 22:52–63.

Smoak, E.M., P. Stein, R.M. Schultz, M.A. Lampson, and B.E. Black. 2016. Long-Term Retention of CENP-A Nucleosomes in Mammalian Oocytes Underpins Transgenerational Inheritance of Centromere Identity. Curr Biol. 26:1110–1116.

Song, K., B. Gronemeyer, W. Lu, E. Eugster, and J.E. Tomkiel. 2002. Mutational analysis of the central centromere targeting domain of human centromere protein C, (CENP-C). Exp Cell Res. 275:81–91.

Stellfox, M.E., I.K. Nardi, C.M. Knippler, and D.R. Foltz. 2016. Differential Binding Partners of the Mis18alpha/beta YIPPEE Domains Regulate Mis18 Complex Recruitment to Centromeres. Cell Rep. 15:2127–2135.

Stoler, S., K. Rogers, S. Weitze, L. Morey, M. Fitzgerald-Hayes, and R.E. Baker. 2007. Scm3, an essential Saccharomyces cerevisiae centromere protein required for G2/M progression and Cse4 localization. Proc Natl Acad Sci U S A. 104:10571–10576.

Sugimoto, K., K. Kuriyama, A. Shibata, and M. Himeno. 1997. Characterization of internal DNA-binding and C-terminal dimerization domains of human centromere/kinetochore autoantigen CENP-C in vitro: role of DNA-binding and self-associating activities in kinetochore organization. Chromosome Res. 5:132–141.

Sullivan, K.F., M. Hechenberger, and K. Masri. 1994. Human CENP-A contains a histone H3 related histone fold domain that is required for targeting to the centromere. J Cell Biol. 127:581–592.

Suzuki, A., B.L. Badger, and E.D. Salmon. 2015. A quantitative description of Ndc80 complex linkage to human kinetochores. Nat Commun. 6:8161.

Suzuki, A., B.L. Badger, X. Wan, J.G. DeLuca, and E.D. Salmon. 2014. The architecture of CCAN proteins creates a structural integrity to resist spindle forces and achieve proper Intrakinetochore stretch. Dev Cell. 30:717–730.

Takeuchi, K., T. Nishino, K. Mayanagi, N. Horikoshi, A. Osakabe, H. Tachiwana, T. Hori, H. Kurumizaka, and T. Fukagawa. 2014. The centromeric nucleosome-like CENP-T-W-S-X complex induces positive supercoils into DNA. Nucleic Acids Res. 42:1644–1655.

Tanaka, K., H.L. Chang, A. Kagami, and Y. Watanabe. 2009. CENP-C functions as a scaffold for effectors with essential kinetochore functions in mitosis and meiosis. Dev Cell. 17:334–343.

Thakur, J., and S. Henikoff. 2016. CENPT bridges adjacent CENPA nucleosomes on young human alpha-satellite dimers. Genome Res. 26:1178–1187.

Tian, T., X. Li, Y. Liu, C. Wang, X. Liu, G. Bi, X. Zhang, X. Yao, Z.H. Zhou, and J. Zang. 2018. Molecular basis for CENP-N recognition of CENP-A nucleosome on the human kinetochore. Cell Res. 28:374–378.

Trazzi, S., G. Perini, R. Bernardoni, M. Zoli, J.C. Reese, A. Musacchio, and G. Della Valle. 2009. The C-terminal domain of CENP-C displays multiple and critical functions for mammalian centromere formation. PLoS One. 4:e5832.

van Hooff, J.J., E. Tromer, L.M. van Wijk, B. Snel, and G.J. Kops. 2017. Evolutionary dynamics of the kinetochore network in eukaryotes as revealed by comparative genomics. EMBO Rep. 18:1559–1571.

Volkov, V.A., P.J. Huis In ’t Veld, M. Dogterom, and A. Musacchio. 2018. Multivalency of NDC80 in the outer kinetochore is essential to track shortening microtubules and generate forces. Elife. 7.

Wan, X., R.P. O’Quinn, H.L. Pierce, A.P. Joglekar, W.E. Gall, J.G. DeLuca, C.W. Carroll, S.T. Liu, T.J. Yen, B.F. McEwen, P.T. Stukenberg, A. Desai, and E.D. Salmon. 2009. Protein architecture of the human kinetochore microtubule attachment site. Cell. 137:672–684.

Watanabe, R., M. Hara, E.I. Okumura, S. Herve, D. Fachinetti, M. Ariyoshi, and T. Fukagawa. 2019. CDK1-mediated CENP-C phosphorylation modulates CENP-A binding and mitotic kinetochore localization. J Cell Biol. 218:4042–4062.

Weir, J.R., A.C. Faesen, K. Klare, A. Petrovic, F. Basilico, J. Fischbock, S. Pentakota, J. Keller, M.E. Pesenti, D. Pan, D. Vogt, S. Wohlgemuth, F. Herzog, and A. Musacchio. 2016. Insights from biochemical reconstitution into the architecture of human kinetochores. Nature. 537:249–253.

Welburn, J.P., M. Vleugel, D. Liu, J.R. Yates, 3rd, M.A. Lampson, T. Fukagawa, and I.M. Cheeseman. 2010. Aurora B phosphorylates spatially distinct targets to differentially regulate the kinetochore-microtubule interface. Mol Cell. 38:383–392.

Xiao, H., F. Wang, J. Wisniewski, A.K. Shaytan, R. Ghirlando, P.C. FitzGerald, Y. Huang, D. Wei, S. Li, D. Landsman, A.R. Panchenko, and C. Wu. 2017. Molecular basis of CENP-C association with the CENP-A nucleosome at yeast centromeres. Genes Dev. 31:1958–1972.

Yan, K., J. Yang, Z. Zhang, S.H. McLaughlin, L. Chang, D. Fasci, A.E. Ehrenhofer-Murray, A.J.R. Heck, and D. Barford. 2019. Structure of the inner kinetochore CCAN complex assembled onto a centromeric nucleosome. Nature. 574:278–282.

Yoda, K., and T. Okazaki. 1997. Site-specific base deletions in human alpha-satellite monomer DNAs are associated with regularly distributed CENP-B boxes. Chromosome Res. 5:207–211.

Zakeri, B., J.O. Fierer, E. Celik, E.C. Chittock, U. Schwarz-Linek, V.T. Moy, and M. Howarth. 2012. Peptide tag forming a rapid covalent bond to a protein, through engineering a bacterial adhesin. Proc Natl Acad Sci U S A. 109:E690–697.

Zasadzinska, E., J. Huang, A.O. Bailey, L.Y. Guo, N.S. Lee, S. Srivastava, K.A. Wong, B.T. French, B.E. Black, and D.R. Foltz. 2018. Inheritance of CENP-A Nucleosomes during DNA Replication Requires HJURP. Dev Cell. 47:348–362 e347.

Zhang, Z., D. Bellini, and D. Barford. 2020. Crystal structure of the Cenp-HIKHead-TW sub-module of the inner kinetochore CCAN complex. Nucleic Acids Res.

